# Using historical museum samples to examine divergent and parallel evolution in the invasive starling

**DOI:** 10.1101/2021.08.22.457241

**Authors:** Katarina C. Stuart, William B. Sherwin, Jeremy J. Austin, Melissa Bateson, Marcel Eens, Matthew C. Brandley, Lee A. Rollins

## Abstract

During the Anthropocene, Earth has experienced unprecedented habitat loss, native species decline, and global climate change. Concurrently, greater globalisation is facilitating species movement, increasing the likelihood of alien species establishment and propagation. There is a great need to understand what influences a species’ ability to persist or perish within a new or changing environment. Examining genes that may be associated with a species’ invasion success or persistence informs invasive species management, assists with native species preservation, and sheds light on important evolutionary mechanisms that occur in novel environments. This approach can be aided by coupling spatial and temporal investigations of evolutionary processes. Here we use the common starling, *Sturnus vulgaris,* to identify parallel and divergent evolutionary change between contemporary native and invasive range samples and their common ancestral population. To do this, we use reduced-representation sequencing of native samples collected recently in north-western Europe and invasive samples from Australia, together with museum specimens sampled in the UK during the mid-19^th^ Century. We found evidence of parallel selection on both continents, possibly resulting from common global selective forces such as exposure to pollutants (e.g. TCDD) and food carbohydrate content. We also identified divergent selection in these populations, which might be related to adaptive changes in response to the novel environment encountered in the introduced Australian range. Interestingly, signatures of selection are equally as common within both invasive and native range contemporary samples. Our results demonstrate the value of including historical samples in genetic studies of invasion and highlight the ongoing and occasionally parallel role of adaptation in both native and invasive ranges.

## 2. Introduction

The ecological and economic impacts of invasive species are a growing concern in our globalised world. Increased intercontinental travel and trade is giving rise to new or reinforced invasion pathways (Turbelin *et al*. 2017), resulting in a great number of alien species becoming established and spreading within novel ranges (Hulme 2009). The financial cost of invasive species within Australia is estimated in excess of 13 billion dollars annually (Hoffmann & Broadhurst 2016). With habitat clearing and climate change expected to favour invasive species over native ones, the environmental and financial cost of invasive species is only expected to rise in the future (Dukes & Mooney 1999). Many studies on invasive species’ success involve examining evolutionary changes following introduction and focus on rapid adaptation to novel environments (Prentis *et al*. 2008). This information is vital for long term management of invasive populations.

Understanding evolutionary trends across a species’ native and invasive ranges will help determine important adaptive elements that aid species’ persistence in a changing world. Species that are invasive present a contrariety when they face population decline within their native range (Rogers *et al*. 2006; Delibes-Mateos *et al*. 2009; Erfmeier & Bruelheide 2010; Bishop 2011). Research efforts should tackle ecological questions of conservation and invasion management concurrently, enabling us to understand how and why patterns of adaptation in a species’ native and invasive populations may differ. It is possible that the translocation and establishment process itself may select for traits that enable an individual to overcome otherwise detrimental environmental instability or other novel stressors, increasing general fitness (Callaway & Ridenour 2004; Liu & Trumble 2007). Understanding how the invasion process may induce differences in population persistence is made even more pressing by the increasing anthropogenic impact on the natural world, including ongoing land alteration, environmental contamination, and human induced climate change (Hellmann *et al*. 2008).

Often, these adaptive changes are identified through contrasting present day native and invasive populations (Hofmeister *et al*. 2021a). However, such approaches exclude the temporal element of species’ change, so that such studies assume native populations have not changed since the founders of the invasive population were collected. This would then lead to the conclusion that all similarities between native and invasive populations result from a common ancestral population and are not due to parallel change since separation. However, with global anthropogenic change impacting the natural world, it is reasonable to assume that altered or increased selective pressures have arisen during the post-industrialised world, shaping species world-wide (Sokolova & Lannig 2008; Siepielski *et al*. 2017). Historical specimens therefore provide an unparalleled tool to better contextualise divergent versus parallel evolution, providing phenotypic and, more recently, genotypic information that can be used to identify temporal changes in species ranges and traits (Ewart *et al*. 2019; Lopez *et al*. 2020). Studies focusing on rapid local adaptation in invasive species may now make use of historic DNA alongside contemporary samples to understand the selective forces shaping both invasive and native ranges concurrently.

The common or European starling, *Sturnus vulgaris*, presents an ideal system to use historical samples to investigate both divergent and parallel genetic change within an invasive species. The European starling (hereafter the starling) is a highly invasive pest, introduced and successfully establishing on every other continent barring Antarctica (Higgins *et al*. 2006). Despite this, the native range starlings are themselves a conservation focus, with declines of more than 50% in some countries (Versluijs *et al*. 2016) putatively associated with shifts in farming practice that are common in their native range (Freeman *et al*. 2007; Heldbjerg *et al*. 2016). Fortunately, due to the historic popularity of collecting bird skins, historical starling samples may be found scattered across many museums and institutions in both their native range and within invaded countries. These skins serve as untapped reserves of genetic information, which may be used to track temporal genetic changes across the native range, reveal information regarding historical population structure, and provide context that enables us to better understand current patterns of native range starling decline.

Starlings present a prime example of how the combination of data from invasive, native, and historical populations can clarify our understanding of evolution in both native and invasive contexts. Introduced into Australia in the 1860’s, the starlings’ range now stretches across the continent’s eastern and southern coasts (Long 1981). Genetic analysis supports strong population sub-structuring across the invasive Australian range (Rollins *et al*. 2009, 2011), with reduced-representation sequencing data indicating the two main subpopulations likely resulted from allelic differences in founding populations at different introduction sites (Stuart & Cardilini et al. 2021a). The historical specimens available for this species were collected within 15 years of the earliest documented introductions to Australia in 1856 (Long 1981), providing a snapshot of native starling populations at the time when founders were transported to Australia.

To better understand patterns of population structure and signatures of selection present in the invasive Australian range, we used a reduced-representation sequencing approach to compare contemporary Australian (AU) and native range (United Kingdom, UK; Belgium, BE) starlings to historical UK samples collected during the period when the Australian founders were collected. Moreover, this project explores proximate drivers of invasive species’ evolution in the face of novel selection provided by new environments. Specifically, we compare population structure of native and invasive contemporary *S. vulgaris* samples, and we explore genomic divergence between contemporary and historical *S. vulgaris* and assess the putatively adaptive capacity of these genomic changes. Finally, we use historic samples as a basis of comparison to determine regions of parallel change in both the contemporary native and invasive populations to better understand global shifts in selective forces.

## 2. Methods

### 2.1 Sample collection and extraction (historical starlings)

We sourced historic starling specimens (HS) from the Natural History Museum (NHM) in Tring, UK (N=15). Historical samples were selected on the basis of sample quality, completeness of collection record, and sample collection date (samples collected from 1857-1871, during the period when the Australian introductions took place; Higgins *et al*. 2006; Supplementary Materials, Table S1).

DNA extractions on historical samples were done in a specialist ancient DNA laboratory at the Australian Centre for Ancient DNA, University of Adelaide. We rehydrated 3-4mm^3^ dried tissue in 1ml of 0.5M EDTA for 2hrs and extracted DNA using a Qiagen DNeasy Tissue Kit (Qiagen, Valencia, California, United States) as per manufacturer’s instructions. DNA was eluted twice with 40 uL of EB buffer (+0.05% Tween-20) for a final elution volume of 80 uL.

### 2.2 Sample collection and extraction (contemporary starlings)

We sourced contemporary native range starling samples from two UK locations: Monks Wood (MW: N=15, blood) and rural sites around the city of Newcastle upon Tyne (NC: N=15, blood), and one Belgian (BE) location in Antwerp (AW: N=15, blood) (Fig. 1a). We sourced contemporary Australian samples from two locations: McLaren Vale in South Australia (MV: N = 15, blood) and Orange in New South Wales (OR: N = 15, muscle tissue) (Fig. 1b). Contemporary DNA extractions were performed using the QIAGEN Gentra Puregene Tissue kit as per manufacturer’s instructions.

**Figure 1:**
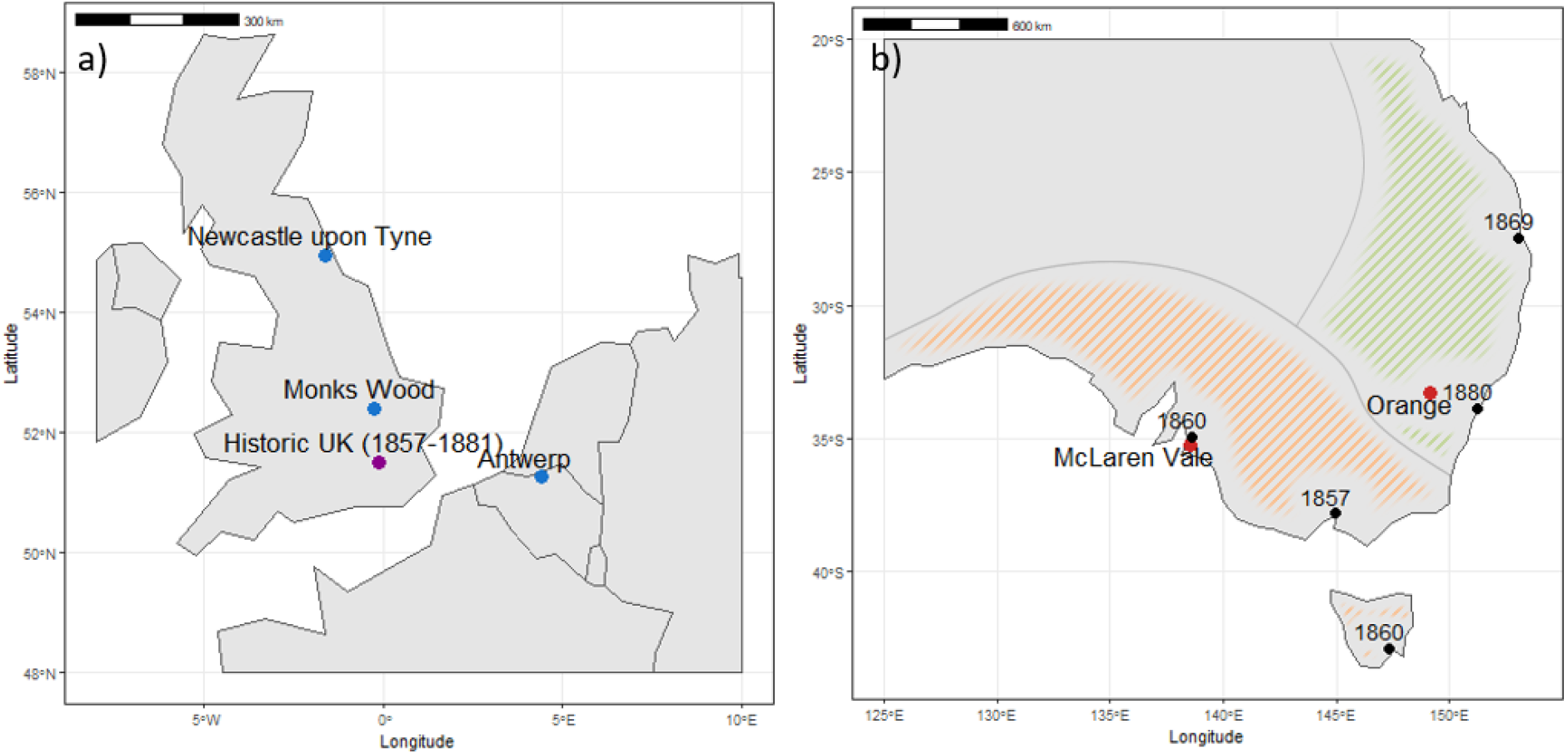
Geographical distribution of the *Sturnus vulgaris* collection sites in the United Kingdom and Belgium (native range, blue and purple), Australia (invasive range, red), and historical samples. The coloured shading on the Australian map denotes their Australian range, broken up into the two main subpopulations. Introduction sites are marked in black on the Australian map, with first introduction year listed adjacently.

### 2.3 Sequencing and genome variant calling

We sequenced 75 contemporary and 15 historical samples using the DArTseq protocol (Kilian *et al*. 2012), using a restriction enzyme double digest of PstI-SphI. The sequencing was conducted on a Hiseq 2500, producing 312,907,523 single-end reads of raw data across the 90 samples (26,408,649 across the 10 successfully sequenced historical samples; Supplementary Materials, Table S1).

We used the Stacks v2.2 (Rochette *et al*. 2019) pipeline (Rochette & Catchen 2017) to process the DArTseq raw data into variant calling format (vcf). We used the *process_radtags* function to clean the tags; discarding reads of low quality (-q), removing reads with uncalled bases (-c), and rescuing barcodes and radtags (-r). We used the Burrows-Wheeler aligner (BWA) v0.7.15 (Li & Durbin 2009) *aln* function to align the read data to the reference genome *S. vulgaris* vAU1.0 (Stuart *et al*. 2021b). Using FastQC, we identified base sequence bias in the adapter region, and so the first five base sequences were trimmed (-B 5) during alignment. We used the BWA *samse* function to produce SAM alignment files, before passing these into Samtools v1.10 (Li *et al*. 2009) to produce BAM files. We ran the cleaned data through STACKS *gstacks* (default parameters) and then *populations* to call the SNPs.

We produced an unfiltered SNP data set by running Stacks *populations* with no parameter thresholds specified. We used this data set to produce the unfiltered loci and site counts (Supplementary Materials, Table S2), split the data into three separate files for further assessment of historic sequencing data (contemporary native range samples: MW, NC, and AW; Australian samples: OR and MV; and historic samples; HS). We calculated variant base substitutions in Vcfstats v0.0.5 (Lindenbaum 2015) and variant density mapped along the reference genome scaffolds using SAMTOOLS *bedtools* function (window size 1,000,000bps). We used the DARTR V1.1.11 (Gruber *et al*. 2018) function *glPlot* to create a smear plot of the mapped variant data across individuals and genomics sites. This resulted in a population genetic file of 239,538 SNPs. We also passed the raw single-end read data through the Stacks BWA-mem, and the bowtie-GATK, variant calling pipeline (Supplementary Material: Appendix 1), to compare the quantity of site variant data that was successfully mapped and assess how these alternate variant calling software performed for reduced representation sequencing data sets that contained degraded historical DNA.

We generated a ‘population genetics’ variant file by running Stacks *populations,* filtering for a minimum per-population site call rate of 50% (-r 0.5), a minimum populations per-site of 2 (-p 2), a minimum loci log likelihood value of -15 (--lnl_lim -15), with one random SNP per locus retained (--write_random_snp). We used VCFTOOLS v0.1.16 (Danecek *et al*. 2011) to filter the following parameters: maximum missingness per site of 50% (-max-missing 0.5), minor allele frequency of 2.5% (MAF; --maf 0.025), minimum loci depth of 5 (--minDP 5), minimum genotype quality score of 15 (--minGQ 15) and site Hardy-Weinberg Equilibrium exact test minimum p value 0.001 (--hwe 0.001). We chose a moderate high threshold for missingness as over-filtering missingness has been found to influence population structure in reduced representation data sets (Wright *et al*. 2019; Selechnik *et al*. 2020). MAF filtering helps remove misreads, and HWE filtering removed highly non-neutral loci, all of which are important for capturing neutral population substructure (HWE SNPs were however left in for admixture analysis (Deng *et al*. 2001)). After filtering, we checked individual relatedness, and closely related individuals were removed so that there was only one representative from each familiar cluster in the final data (Supplementary Materials: Fig. S1). This resulted in a population genetic variant file of *23,683* SNPs using in the below section ‘Population structure analysis’ (*25,572* variant sites if HWE were included).

We generated a ‘selection’ variant file by using Stacks *populations* to align the raw reads (with --lnl_lim -15 flag), and then used vcftools to filter out only reads present in at least 50% of the historic individuals (i.e. in at least 5 historic individuals), with additional quality filtering (--minGQ 15 --minDP 2), resulting in 12,219 SNP sites. Only these sites were then retained to filter the original *populations* variant file, along with a MAF minimum of 2.5% to remove possible sequencing errors. This produced a data set with fewer SNPs than the population genetics variant file but retained only SNPs sequenced in a moderate number of historic individuals, which would be necessary for selection analysis. We filtered the selection variant file to form three pairwise population SNP data comparisons; UK-HS (UK populations MW, NC, BE and 10 historic individuals), AU-HS (AU populations MV and OR, and 10 historic individuals), and UK-AU (UK populations MW and NC, and AU populations MV and OR). While the native range population may contain a mix of resident and migratory individuals, because we see minimal population structure in the native range and very small Fsr values (0.003-0.008) we decided to include all contemporary native range samples in this analysis. Within each of these three variant file subsets, we retained SNP sites present in at least 5 contemporary individuals, as the file was already filtered for loci present in at least 50% (5/10) of historic individuals. This relatively lenient filtering was necessitated by the smaller number genomic sample sites produced by the degraded DNA, and ensured minimal loss of the historic information that was present. This resulted in three pairwise population files used in the below section ‘Selection analysis’: UK-HS (*4997* SNPs), AU-HS (*5006* SNPs), and UK-AU (*4957* SNPs).

### 2.4 Population structure analysis

We analysed the population genetics variant file several ways to examine the population structure and differentiation between and within the historical and contemporary sample regions. We used R v

3.5.3 (R Core Team 2017) to run the SNPRelate *snpgdsPCA* function to create a principal components analysis (PCA) of the loci. We used Admixture v1.3.0 to determine individual ancestry proportions for each of the following three sample subsets: all samples, contemporary native range and historic, and contemporary Australian. We calculated marginal likelihood for model complexity (K) 1-8 by averaging over 50 runs, and admixture proportion (Q) profile were generated by Clumpak (Kopelman *et al*. 2015) (run on default settings) to obtain an average Q profile from 25 runs. We used the dartR function *gl.dist.pop* to calculate the Euclidean distance between sampling locations, and the STAMPP function *stamppFst* to calculate pairwise FST between sampling locations (nboots=1, percent=95, nclusters=1). Finally, we used the dartR function *gl.tree.nj* to visualise phylogeny of the 6 sampling groups.

### 2.5 Selection analysis

When looking for diversifying selection, allele frequency based approaches are often used, such as Bayescan v2.1 (Foll & Gaggiotti 2008). Bayescan aims to identify SNPs subject to natural selection by assigning a per-site posterior probability estimated by comparison of explanatory models with and without selection. We conducted Bayescan SNP outlier analysis, with prior odds for the neutral model set to 10 (-pr_odds 10), and a false discovery rate (FDR) of 0.05.

Because the smaller sample sizes in this selection analysis lack statistical power, and Bayescan uses stringent criteria to flag selection, the alpha values produced by Bayescan were used to identify additional putative sites under selection. The alpha value is the locus-specific component of selection identified in the Bayescan selection model, where a large positive alpha value indicates diversifying selection, and a large negative value indicates balancing or purifying selection. We retained sites that reported an alpha value of >0.1, and then filtered these for a minimum r2 linkage value of 0.05 (window size 1000bp) using SAMTOOLS *bcftools*, similar to the approach used in Gloria-Soria et al. (Gloria-Soria *et al*. 2016). We plotted Alpha values in rank order and chose an alpha threshold of 0.1 as it represented a reasonably stringent cut-off across all three pairwise comparisons (see Supplementary Materials: Fig. S3). Using the less stringent means of alpha value to categorise a SNP as under putative selection will yield more false positives, hence we chose to filter for high linkage. Filtering out sites with low linkage will help select against false positive sites whose allele frequencies (and hence alpha value) appear to indicate selection, because linkage often occurs around a gene under selection (Slatkin 2008).

Degraded historical DNA from older museum skins is known to have bias towards low diversity SNPs (Ewart *et al*. 2019), which may impact single site approaches to examining divergent selection. Therefore, we used a F_ST_ sliding window approach to identify genomic regions of putative diversifying selection. VCFTOOLS *weir-fst-pop* function was used to analyse weighted F_ST_ in 900000 bp windows (10000 window step). We chose this window size primarily based on the ratio of variants to genome size, with the chosen windows putatively spanning 3-4 variants, with small step sizes then allowing shifts in F_ST_ patterns to be pinpointed more exactly. We selected site windows that reported a weight F_ST_ value in the top 99th percentile for each pairwise population selection analysis and analysed the Fst of SNPs within these windows in rank order. We retained any SNP within these outlier windows that lay above an Fst threshold relevant for each pairwise data set (this was determined visually as a plateauing of the ranked Fst values, see Supplementary Materials: Fig. S4) as putative outlier SNPs.

From each of the three above identification processes, we pooled putative outlier SNPs within each pairwise population comparison, to be used for further variant analysis.

### 2.5 Variant analysis and annotation

SNPs that were reported as outliers across either the UK-HS or AU-HS data set, as well as the UK-AU data set, we designated as sites under divergent selection. SNPs that were reported as outlier across both UK-HS and AU-HS data sets we designated as sites under parallel selection. The remaining outlier SNPs we identified only in one dataset: UK-HS (putative UK selection); AU-HS (putative AU selection); or UK-AU (putative UK-AU divergence). Further details are given below in section 3.4 Putative adaptive selection.

We analysed these five groups of SNPs for their functional roles and the nature of the mutation. We completed SNP analyses primarily using Variant Effect Predictor (VEP) (McLaren *et al*. 2016), using the genome annotation version released alongside the *S. vulgaris* vAU1.0 assembly (Stuart *et al*. 2021b) to examine the functional consequences of the SNPs (processed to exclude multiple isoforms using AGAT *agat_sp_keep_longest_isoform.pl* (Dainat 2020)). We used Bedtools and the Agat function *agat_sp_functional_statistics* to extract genes and transcripts that overlapped the putative loci under selection, and extract gene ontology (GO) terms. We used Revigo (Supek *et al*. 2011) to visually summarise GO terms, and we calculated allele frequencies at SNP sites using bedtools.

Lastly, to test if there was an overrepresentation of SNPs located on the macrochromosomes (>20 Mb, as described in Backström et al. 2010), microchromosomes, or the sex chromosome, we used a Chi-square test to examine overrepresentation of these SNP types across four different SNP groupings: the divergent SNPs (both UK and AU), the parallel SNPs, the SNPs under selection (Putative UK, AU, and UK and AU SNPs under selection), and the remainder of the SNPs that were not flagged as putatively being under selection in any of the pairwise data sets (no selection). We analysed these data using the *chisq.test()* function in R. To ensure results from this analysis were not artifacts of the data (for example due to diploid variant calling on hemizygous ZW females), we conducted two types of analysis of variance (ANOVA) tests. Firstly, we tested to see whether major allele frequency was significantly associated with an interaction between SNPs type (selection vs non-selection) and SNP location (sex chromosome vs autosome). Secondly, we examined whether total number of alleles sequenced for a SNP site was significantly associated with an interaction between SNPs type (selection vs non-selection) and SNP location (sex chromosome vs autosome). We tested for an interaction between SNPs type (selection vs non-selection) and SNP location (sex chromosome vs autosome on either the major allele frequency or number of alleles sequenced for a SNP site using the *aov()* function in R. We constructed this analysis as both major allele frequency and allele counts are employed to flag outliers to the outlier tests used in this study, and an interaction between SNP location and its categorisation as under selection or not would be cause for concern about bias due to data artifacts. This analysis was run with the major allele frequency and allele counts of all individuals, and then separately again for just historic individuals, so see if any of these effects were only affecting historic individuals (e.g. due to smaller sample size, which may make historic allele frequencies more vulnerable to sex chromosome biases).

## 3. Results

### 3.1 Population structure of contemporary and historical *S. vulgaris*

Our genetic data revealed strong differentiation between contemporary native and invasive range samples, as well as replicating the previously established subpopulation structure within Australia. Using PCA, the historical samples were genetically intermediate between the contemporary UK and AU populations along the first PC axis (Fig. 2a), though undoubtedly part of this central placement was owing to higher levels of missing data in historic individuals due to degraded DNA (Supplementary Materials: Table S2).

**Figure 2:**
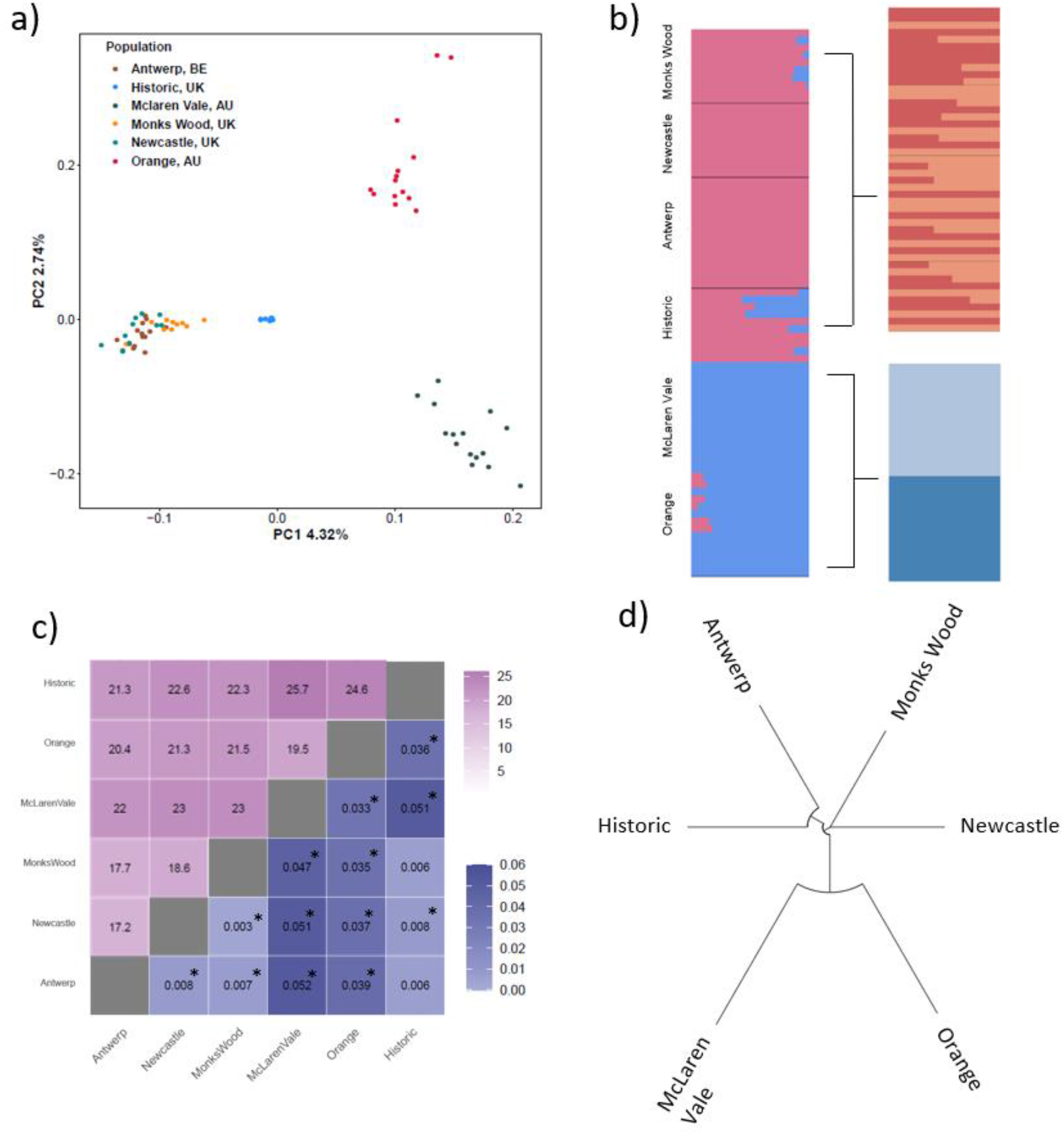
**Population genetic analysis for contemporary and historical *Sturnus vulgaris*** using the population genetics variant file. Panel a) denotes a PCA of the 6 sampling groups using SNPRELATE, panel b) denotes ADMIXTURE ancestry Q profiles, averaged over 50 runs using CLUMPAK for all sample groups, all native range samples (contemporary and historic), contemporary native range samples, and invasive Australian samples. Panel c) displays a heatmap of pairwise analysis between each of the sample groups, with above the diagonal (purple) indicating Euclidean distance, and below the diagonal (blue) indicating pairwise F_ST_ (asterisk * denoting a significant F_ST_ result), and panel d) displays the phylogenetic relationships between sampling sites using a neighbour-joining tree.

Through admixture analysis on the 85 sequenced samples (five historic samples failed sequencing, see section 3.4 Sequencing and variant calling with historic samples) we determined K=1 and K=2 had similar support (Supplementary Materials: Fig. S2a). Ancestry proportions when plotted for K=2 (Fig. 2b) support the historic UK clustering closer to the contemporary native range samples (though containing a mixture of native [red] and invasive [blue] genotypes), with very little further substructure revealed when admixture analysis was conducted just on the native range samples (Fig. 2b, Supplementary Materials: Fig. S2b&c). We identified strong substructure in the Australian samples (Supplementary Materials: Fig. S2c).

Of the invasive Australian samples, we found that those from McLaren Vale showed greater genetic differentiation from the native range samples than did those from Orange, evident in pairwise F_ST_ and Euclidean genetic distance comparisons (Fig. 2c) and corroborated by McLaren Vale admixture proportions lacking the UK cluster (Fig. 2b). The historical samples were found to be most similar to samples from Antwerp, when considering pairwise F_ST_ and Euclidean genetic distance (Fig. 2b) and tree analysis (Fig. 2d), while PCA and admixture plots are indicative of more similarity between historical samples and those from Monks Wood. We also characterised the relationship amongst the native range samples by the neighbourhood-joining tree (Fig. 2d). We found that the genetic differentiation between sampling sites across the native range is less than that across the invasive range.

### 3.2 Genomic divergence between contemporary and historical *S. vulgaris*

Using the default Bayescan pipeline, we identified a total of 24 outlier loci across the three pairwise comparisons, fifteen of which were found in the UK-AU comparison, eight in the UK-HS comparison, and only one in the AU-HS comparison (Table 1, Supplementary Materials: Fig. S3a-c). Using the Bayescan alpha+LD approach, we found a roughly similar amount of outlier sites across both the UK-HS and AU-HS, with approximately half as many identified as significantly different between UK-AU (Table 1, Supplementary Materials: Fig. S3d-f). Interestingly, we found that across the three pairwise population analyses, not all of the outlier SNPs reported by the Bayescan FDR approach were shared with the Bayescan alpha+LD approach (Supplementary Materials, Table S3), presumably because they did not meet the linkage criteria. Using the sliding window approach, the AU-HS comparison reported marginally the highest number of putative outlier loci compared to UK-HS and UK-AU (Table 1, Supplementary Materials: Fig. S4). Most but not all the SNPs identified by the Bayescan FDR approach we also identified in the F_ST_ sliding window approach, with generally high overlap with the Bayescan alpha+LD approach (Supplementary Materials, Table S3). We visualised the outlier sites across all three methods against the starling genome for each of the pairwise population data sets, and they appear to be fairly uniformly spread throughout the genome (Fig. 4).

**Figure 4:**
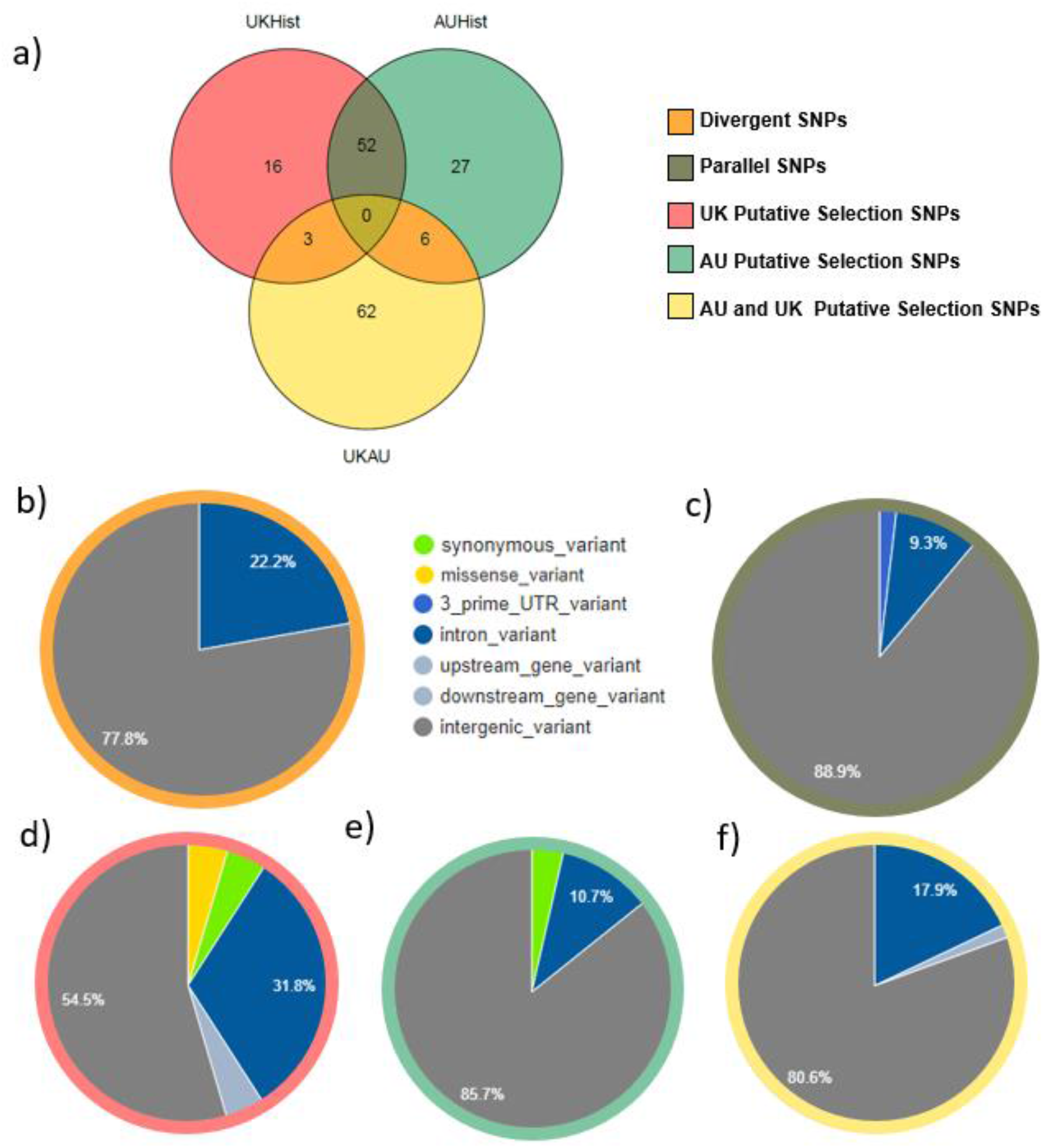
Summary of *Sturnus vulgaris* putative SNPs under selection, with panel **a)** depicting a Venn diagram of group categorisation, and the remaining figures the Variant Effect Predictor (VEP) summary outputs of function variation for each of the five SNP groups of **b)** SNPs under divergent selection, **c)** SNPs under parallel selection, **d)** UK SNPs under putative selection, **e)** AU SNPs under putative selection, and **f)** UK and AU SNPs under putative divergence.

**Table 1:** Number of putative sites under selection in *Sturnus vulgaris* reported by the different selective scans for the pairwise comparisons of UK-HS, AU-HS, and UK-AU data set.

| Outlier group |  | UK-HS | AU-HS | UK-AU |
| --- | --- | --- | --- | --- |
| data set SNP count |  | 4997 | 5006 | 4957 |
| Bayescan | *FDR 0.05* | **8** | **1** | **15** |
|  | *Alpha >0.05 + LD*<br>*r<sup>2</sup> >0.5* | **62** | **69** | **34** |
| Fst Sliding Windows | *Fst > 0.5 + SNP*<br>*threshold* | **27** | **37** | **33** |
| Total |  | 71 | 85 | 71 |

We pooled putative outlier SNPs across all the analyses done for each pairwise comparison, yielding a total of 71, 85, and 71 unique SNPs across identification methods for UK-HS, AU-HS, and UK-AU respectively (Table 1).

### 3.3 Putative adaptive selection

Of the total of 227 SNPs identified as under selection across all three data sets, we found 166 unique SNPs, of which 9 were identified as resulting from divergent selection, and 52 from parallel selection (Fig. 4a). The remaining SNPs were only present in one of the pairwise data sets, and were categorised as resulting from putative UK selection (16), putative AU selection (27) and putative UK and AU divergent selection (62) (Table 2, Fig. 4a).

**Table 2:**
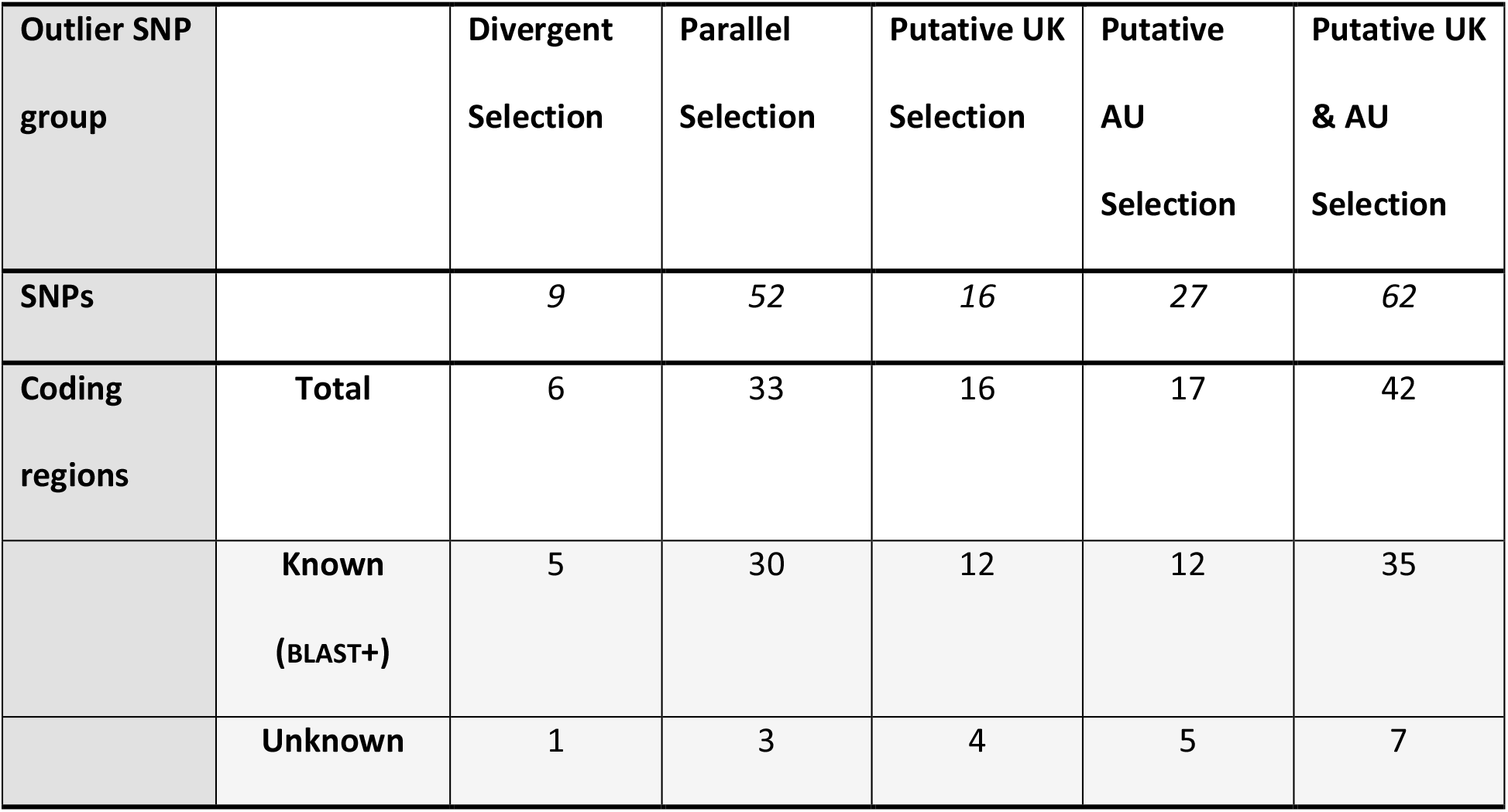
SNP and gene counts of *Sturnus vulgaris* SNPs putatively under selection, for the five categorical grouping based on pairwise comparisons.

| Outlier SNP group |  | Divergent Selection | Parallel Selection | Putative UK Selection | Putative AU Selection | Putative UK & AU Selection |
| --- | --- | --- | --- | --- | --- | --- |
| SNPs |  | 9 | 52 | 16 | 27 | 62 |
| Coding regions | Total | 6 | 33 | 16 | 17 | 42 |
|  | Known (BLAST+) | 5 | 30 | 12 | 12 | 35 |
|  | Unknown | 1 | 3 | 4 | 5 | 7 |

Across all five of these data sets, VEP analysis of the functional nature of the SNPs revealed that variants were mostly intergenic, and there were some within-intron variants (Fig. 4b-f). Of the SNPs under divergent and parallel selection, 80% were intergenic variants, with the remained largely being made up of intron variants, and one 3’ UTR variant being flagged as under parallel selection. SNPs identified as under UK putative selection (UK-HS divergence) contained the fewest proportion of intergenic variants (Fig. 4d). We found only a few SNPs with predicted protein coding consequences, with one missense variant present in the UK putative selection SNP list, and one synonymous mutation each in the putative UK selection and putative AU selection SNPs (Fig. 4c and 4d).

While many of the SNPs mapped to unannotated loci, we mapped five SNPs in the divergent data set to annotated genes, along with 26 SNPs in the parallel data set (Table 2, Supplementary Materials; Table S4, Table S5). Only one gene was found to be divergent within the native range (*ANKHD1*), while four were found to be diverging with the invasive Australian range (*GRIK2*, *Esrrg*, *ARHGAP10*, and *Cacna2d3,* Supplementary Materials; Table S5). A diverse range of genes were flagged as being under divergent parallel selection in both the native and invasive range, including two that are known to interact with toxins and pollutants (*NID2* and *SOS1),* and two related to carbohydrate processing (*SI* and *INS,* Supplementary Materials; Table S5). There was little gene ontology overlap between these two data sets (Supplementary Materials; Fig. S5).

Using the Chi-squared analysis, we found that within SNPs under putative selection there was not a proportional distribution of SNPs across macro, micro, and sex chromosomes, (χ^2^ (6, N = 5068) = 21.149, p = 0.0017). We visualised the residuals and contributions, revealing an overabundance of sex chromosome SNPs for both the parallel SNPs as well as the UK, AU and UK and AU SNPs under putative selection (i.e. those SNPs that were reported as an outlier in only one pairwise population comparison; Fig. 6). We conducted additional analysis on potential biases in major allele frequency or allele counts to help validate this result. Across all samples, we found no significant interaction between SNP location (sex chromosome or autosome) and SNP category (under selection or not) for either major allele frequency (F_1,5064_ = 0.188, p-value = 0.664) or allele counts (F_1,5064_ = 0.278, p-value = 0.598). We repeated these analyses for historic individuals only, which likewise found no significant interaction between SNP location and SNP category for either major allele frequency (F_1,5064_ = 1.757, p-value = 0.185) or allele counts (F_1,5064_ = 0.668, p-value = 0.4136).

**Figure 6:**
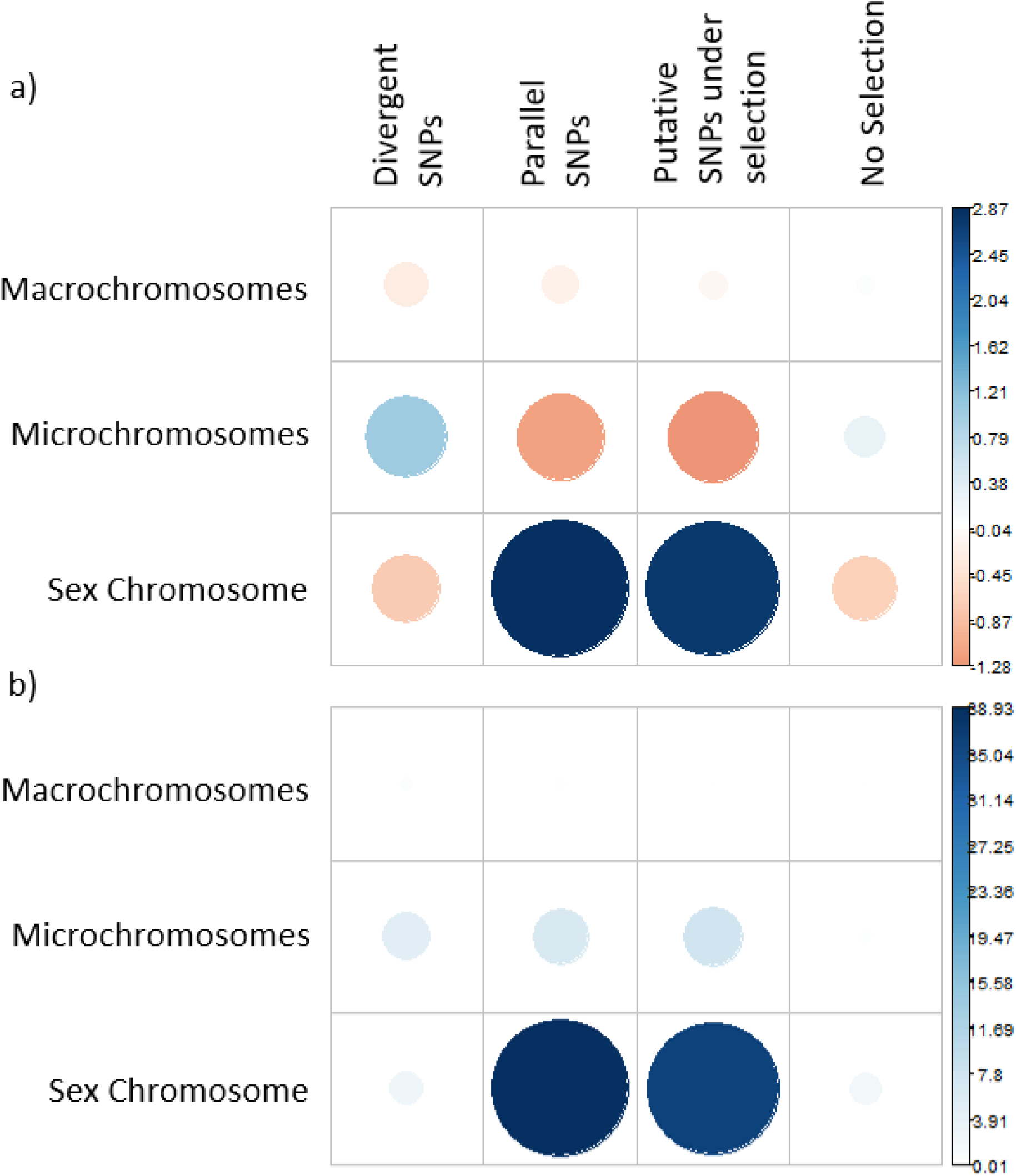
**Test of statistical association between SNPs categorised as under selection versus the chromosome type they reside in for *Sturnus vulgaris* DArT-Seq**, where panel a) depicts visualization of Pearson residuals, where the size of the circle is proportional to the amount of the cell contribution, positive residuals (indicating a positive correlation) are in blue, and negative residuals (indicating a negative correlation) are in orange, and panel b) depicts relative contribution of each cell to the total Chi-square score.

### 3.4 Sequencing and variant calling with historic samples

Of the 15 historical samples, 10 were successfully sequenced using DArTseq, a success rate (66.7%; Supplementary Materials, Table S1) similar to that previous reported (62%) in a study using museum toe-pad samples ranging from 5-123 yrs old (Ewart *et al*. 2019). There did not appear to be any trends related to DNA concentration, sample age, or fragmentation between historical tissue samples that were successfully sequenced and those which were not (Supplementary Materials: Table S1; Fig. S6, Fig. S7).

Of the three variant calling pipelines (Supplementary Materials, Table S2), we found that Bowtie2-GATK reported the highest percentage of successfully mapped reads over both contemporary and historical samples, followed closely by BWA-Mem with BWA-Aln having much lower mapped read percentages. However, Bowtie2-GATK reported the smallest numbers of variant sites in the unfiltered and filtered data set, with BWA-Mem reporting slightly higher values than BWA-Aln. These results are in alignment with previous assessments of these software performances for reads of approximately this length (Li & Durbin 2009; Li 2013). BWA-aln is generally reported to map more conservatively than BWA-mem (Robinson *et al*. 2017), leading to the much smaller mapped reads percentage, but this did not have a very large effect on the site counts or missing data per individual. The biggest difference between these two was the difference between the number of filtered variant sites for the historical samples, indicating that the lower quality reads produced by the historical samples were most impacted by the change in aligning algorithm. BWA-Aln was used as the variant calling pipeline in this paper because our read length fell on the border of what was recommended for BWA-Aln and BWA-Mem (70 BP), and a more conservative mapping and variant calling approach is suitable for population and selection analysis (when approaches are based on per-site allele frequencies).

We assessed the unfiltered data, and the base substitution plots per-population revealed that though the historical samples reported lower SNP counts, base substitution frequencies were similar across the three population groupings (Supplementary Materials: Fig. S8a-c). When we mapped and aligned these reads to the genome assembly alongside Illumina whole genome variant data for the species (Hofmeister *et al*. 2021a), similar patterns are seen between the two sequencing approaches, and across all three population groupings, though with lower resolution in the historical individuals (Fig. 5). Our sequencing and mapping of the historical samples indicate that, despite the lower quality and fragmented DNA, the overall patterns of base substitutions resembled that of the higher quality fresh tissue used from the contemporary populations, and that the reduced representation approach reflected variant densities seen in whole genome sequencing analyses. A smear plot of data revealed that missing data are relatively evenly spaced along the genome for historical samples (Supplementary Materials: Fig. S9), and not centred on particular genomic regions or chromosomes. Finally, the MAF plots revealed slightly differing patterns of minor allele frequency across the six sample groupings sequenced but, importantly, the historical samples did not appear to contain an unusually high number of sites with very low MAF (Supplementary Materials: Fig. S10)

**Figure 5:**
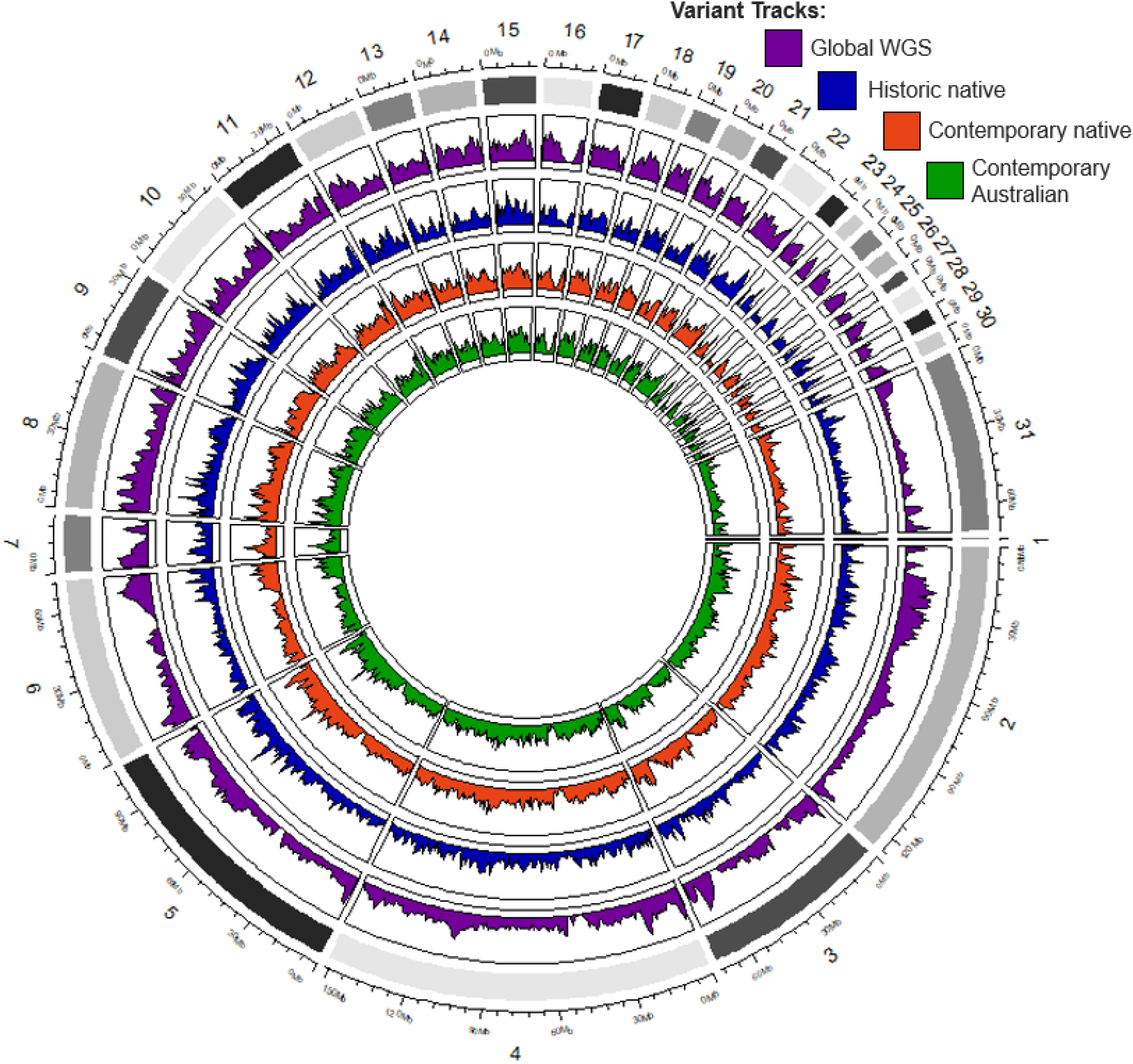
***Sturnus vulgaris* WGS and reduced representation sequencing SNP variant density** plotted in 1,000,000 bp windows around the genome (first 31 scaffolds) for Illumina whole genome global starling variant data (purple), historical samples (blue), contemporary native range samples (red) and contemporary Australian samples (green).

## 4. Discussion

This study demonstrates that *S. vulgaris* has not only undergone divergent selection within the invasive Australian range, but that the native and invasive ranges are undergoing parallel selection, possibly in response to global environmental changes. We note that contemporary native range populations, when considering comparative numbers of divergent SNPs, have undergone a similar amount of genetic change when compared to invasive range populations, despite the latter presumably being exposed to radically different and novel selection regimes. Moreover, we identified several genes related to pollution and carbohydrate metabolism that appear to be under parallel selection in the contemporary native and invasive range samples, which may reflect global environmental changes over the last century and a half. The genes reported as divergent between the populations capture differing selection regimes driving evolution within the two contemporary populations. We also identified a bias for selection on the Z chromosome in comparison to the autosomes. Importantly, this study has successfully used reduced representation sequencing of historic and contemporary specimens to examine selection in *S. vulgaris* within the native and Australian invasive ranges. While the success rate and quality of the historic specimen sequencing reads was less than the contemporary counterparts, the method nevertheless yielded sufficient SNP data to enable examination of population structure and description of temporal patterns of genomic change in starling populations.

### 4.1 Population structure

Very little native range *S. vulgaris* genetic data exists, hence our study provides much needed insight into the population structure and genetic variation of the UK region of the native range. We identified low levels of genetic differentiation across the native range. Some native range starlings are migratory (Feare 1984), and this large-scale dispersion undoubtedly helps to maintain genetic diversity and suppress local differentiation. As expected, the historical starlings bear a stronger genetic resemblance to contemporary samples from the native range than those from the invasive Australian population. The historical samples are most differentiated from their contemporary counterparts in the PCA analysis as compared to analyses of admixture, F_ST_, genetic distance and phylogeny. Interestingly, admixture analysis depicts historic samples as a mixture of contemporary native and invasive genotypes, though with the UK cluster appearing more predominantly. Genotype substructure is therefore indicative of shifting starling genetics over the last 160 years within the native range.

Records identify that the historical samples were taken from around London. Given this, it is interesting that the only contemporary UK population that had a statistically different F_ST_ measure from historic populations was Newcastle. However, it should be noted that while these Fst values are statistically different, they are not biologically important as they represent a negligible difference in allelic variance that may have been affected by study specifics such as sampling scheme. The different population genetics analyses conducted reported that either Monks Wood or Antwerp bears the strongest resemblance to the historic samples, the different results likely a result of different underlying statistical approaches between alternative analytical approaches (e.g. using allelic frequencies for F_ST_, and ordination dimension reduction for PCA). The genetic differentiation between the two invasive Australian population concurs with the two previously described Australian genetic subclusters (Stuart & Cardilini et al. 2021a), and further reinforces the idea that there were slight but distinct genetic differences in the disparate introductions of founding individuals. Comparing the contemporary Australian sample sites to GBS sequencing in the same regions (Stuart

& Cardilini *et al*. 2021a), suggests that the sample sizes in this study were sufficient to be representative of the genetic variation at sampling locations, and that between sample site genetic divergence is higher within this invasive when compared to the native UK over a comparable geographic distance (Stuart & Cardilini *et al*. 2021a).

### 4.2 Genomic divergence

Of the three outlier selection methods used, the default Bayescan approach identified the fewest sites. This was likely due to the small sample sizes lending less statistical power to this stricter analysis, meaning the program was unable to pick up the low signals of selection in these recently diverged populations (Al-Breiki *et al*. 2018). Interestingly, SNPs that were designated as outlier SNPs using Bayescan approaches did not always align with F_ST_ peaks from what, particularly in the UK-AU data set (Fig 3). These flagged outliers with lower SNP site F_ST_ were made up proportionally of the SNPs flagged using the Bayescan default and alpha+LD approach (with the Fst outlier method inherently selecting against lower SNP site F_ST_). Because the UK-AU data set contained a larger number of individuals than the pairwise comparisons that incorporated the historical data, it is reasonable to conclude that these are likely legitimate but weaker outliers that were able to be identified due to increased statistical power. With smaller historical sample sizes, detection will likely be biased towards highly diverged SNPs, which have high F_ST_ values.

**Figure 3:**
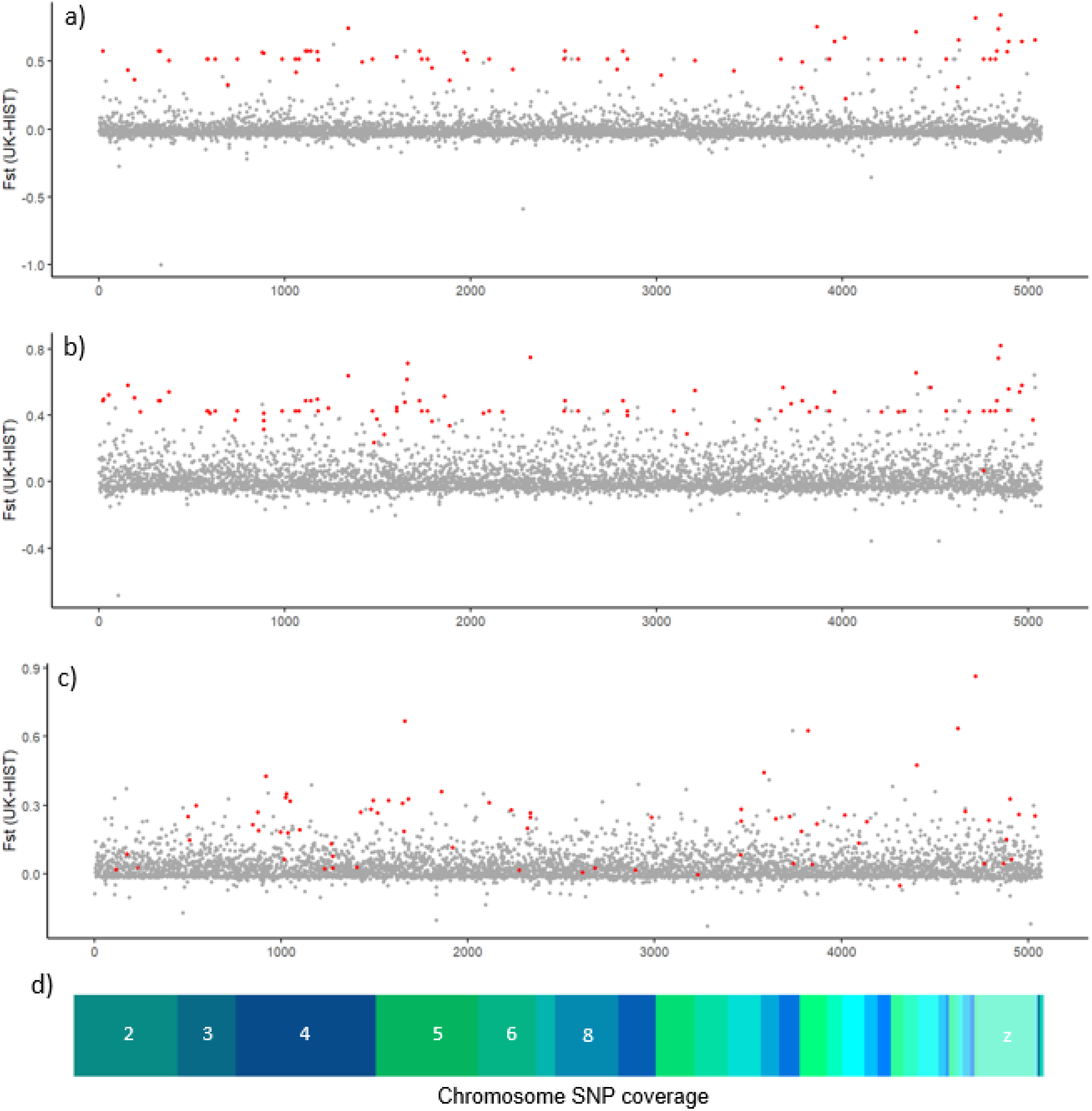
Mapping putatively selected loci across the *Sturnus vulgaris* genome. Panels **a)** (UK-HS samples), panel b) (AU-HS samples), and panel **c)** (UK-AU samples) display the weighted FST in 900kb (10kb step) sliding windows as calculated by vcftools. Non-outlier SNPs are plotted in grey, while outlier SNPs are plotted in red. Panel **d)** depicts the major scaffolds of the *S. vulgaris* genome assembly, sized for their representation in the SNPs plotted in panels a-c).

We identified a larger number of SNPs categorised as being under putative UK & AU selection (62 SNPs), in comparison to putative UK selection (16) and putative AU selection (27). This is likely because lower statistical power of smaller historic sample sizes failed to flag some sites under relatively weaker selection in the UK-HS and AU-HS pairwise analysis. These aforementioned SNPs were the ones that were only flagged in one of the pairwise comparisons, and therefore were not categorised as either parallel or divergent. Interestingly, for the SNPs that were categorised, we see approximately five times the number of SNPs under parallel selection (52 SNPs) in comparison to divergent SNPs (9 SNPs). Across these five SNP selection lists, we identified few variants that were predicted to alter coding potentials within a gene. Other than intergenic variants, intron variants made up the highest proportion of SNPs. Despite not being transcribed gene regions, introns may function as gene regulatory regions, so polymorphisms may still eliciting functional changes (Shaul 2017). We also observed a bias towards the larger sex chromosome (Z) in terms of SNPs that were categorised as being under parallel selection (and also within the UK, AU, and UK and AU SNPs under putative selection). It is possible that this may have arisen due to biases in aspects of the data (e.g. sequencing method, SNP variant calling pipeline). However, we found no differences in major allele frequency or allele variant numbers (factors which may impact a SNP site’s ability to be flagged in the outlier analysis methods used) across SNPs categorised as either under selection or not, across sex chromosomes and autosomes. The conclusion does align with theory that suggests that sex chromosomes are capable of playing a disproportionate role in evolutionary divergence due to their haploid nature in one sex (e.g. the ‘faster-X effect’; Meisel and Connallon 2013), and may be one of the first steps towards speciation (Oyler-McCance *et al*. 2015; Wilson Sayres 2018).

SNPs under putative parallel selection were mostly seen to have higher allelic diversity in the historical samples, with both contemporary populations becoming fixed (or nearly so) for the same allelic variant (supplementary materials, Table S4). This may be indicative of beneficial/non-deleterious alleles shifting towards fixation within contemporary populations, linkage with a nearby variant, or be a result of random processes such as drift (though this latter explanation would be quite unlikely for parallel processes). This over-representation of fixation in contemporary populations is unsurprising, because parallel evolution is more likely to have been based on standing genetic variation in the ancestral population (variants that were already there) than on the same novel variants arising independently in the two descendant populations. We do see apparent fixation in historical populations and more mixed allele frequencies within contemporary populations, in a small number of cases within the parallel SNP data. Divergence within the AU population may be a result of invasion bottleneck processes biasing allelic variation for or against rare variants (depending on their representation in the translocated individuals). However, given that several hundred individuals were introduced to Australia, it would be very unlikely for common allelic variants to be lost through such random processes and more likely that the observed allele frequencies are a result of factors post-introduction (e.g. selection against non-local maladaptation, drift).

### 4.3 Genes undergoing putative adaptive selection

Of the genes that have possibly undergone parallel selection in both the native and invasive range, we see a number of these genes which have been previously associated with responses to various pollutants. Exposure to HT-2, a grain mycotoxin, has been shown to downregulate *nidogen 2* (*NID2*) expression in bovines (Li *et al*. 2020a). Starling interaction with livestock is well documented, and indeed is a key motivating factor of their management in agricultural areas (Linz *et al*. 2007). With increasing livestock numbers, grazing areas, and feed grain use (Spragg 2018), it is not unreasonable for birds that opportunistically feed from grain stocks to face increased exposure to grain-borne toxins and pollutants. Another pollutant, TCDD (2,3,7,8-tetrachlorodibenzodioxin), is persistent and globally used, and has been documented to alter cell cycle progression and proliferation, mediated through the gene *Son of sevenless homolog 1* (*SOS1*) (Pierre *et al*. 2011). Dioxins have a high solubility in lipids, and coupled with a chemical half-life of 9-100 years (depending on substrate), bioaccumulation of these toxins in animals and plants is a well-documented and significant health concern (National Dioxins Program (Australia) *et al*. 2004). Over the last 160 years, increased modernisation globally has resulted in organisms being exposed to increased levels of pollutants. Even trace amounts of compounds may be detrimental to some organisms (Kozlov *et al*. 2009; Bucci *et al*. 2020), and starling eggs have been demonstrated to accumulate polluting organic compounds (Eens *et al*. 2013). Global bird numbers have been declining, across both rare and common species (Gross 2015; Li *et al*. 2020b), with particular toxins such as neonicotinoid from insecticides being documented to play a major role in avian insectivore decline (Hallmann *et al*. 2014). Understanding which pollutants species are coping with and which are not may be important for understanding species and range persistence patterns and may shed light onto the various factors influencing native range starling declines.

Two genes related to carbohydrate metabolic processing were flagged as under parallel selection across the native and invasive range: s*ucrase-isomaltase, intestinal* (*SI*) and *insulin* (*INS*). Gene *SI* encodes for an intestinal tract enzyme involved in breaking down sucrose (obtained from e.g. fruit) and maltose (e.g. grain). Further, *INS* is a pivotal hormone that regulates the metabolism of carbohydrates and lipids. Starlings naturally feed on invertebrates, however shifts towards intensive animal farming and agricultural land expansion globally (Macdonald & McBride 2009) have likely caused increased opportunistic feeding on sucrose and maltose via livestock feed (Linz *et al*. 2007). While evidence suggests that starlings possess limited sucrose processing ability (Martínez del Rio 1990), alterations of allele frequencies within these two genes in both AU and UK populations has implications for selection on metabolic processes that may interact with human supplied food sources, either in the face of the increasing availability of human-supplied food or due to declines in invertebrate numbers (Hallmann *et al*. 2017).

Within the genes that are possibly undergoing divergent selection across the native and invasive range, gene ontology analysis indicates that the biological processes of signal transduction and ion transport are over-represented. Only one gene was reported as undergoing divergence within the native range: *Ankyrin repeat and KH domain-containing protein 1* (*ANKHD1*) (supplementary materials, Table S4, Table S5). ANKHD1 plays an important role in cell cycle progression and proliferation, and has been associated with cancers in humans and model organisms (Dhyani et al. 2012; Machado-Neto et al. 2014). Four genes, *estrogen related receptor gamma* (*Esrrg*), *Glutamate Ionotropic Receptor Kainate Type Subunit 2* (*GRIK2*), *Rho GTPase-activating protein 10* (*arhgap10*), and *Voltage-dependent calcium channel subunit alpha-2/delta-3* (*Cacna2d3*) are reported as divergent in the invasive AU range (supplementary materials, Table S4, Table S5). Of particular interest, the gene *Esrrg* is important for regulating metabolism and energy production in cells, and has been identified as a possible adaptive feature of the desert environment in dromedary camels (Bahbahani *et al*. 2019). It is feasible that all these biological functions would assist starlings as they established and colonised the novel and arid environment of Australia. Doubtless, comparisons to starlings sampled from other arid invasive regions (inland North America and South Africa), as well as the northern African native range edges would provide insight into arid adaptation variation in this species globally.

Of the remaining genes under divergent selection in AU, *ARHGAP10* is involved with Rho signalling-mediated neurogenesis, and may be a regulatory gene related to flightlessness in Galapagos cormorants (Berger & Bejerano 2017). Lastly *GRIK2,* involved in the functional molecular organisation of the avian cerebrum (Jarvis *et al*. 2013), and *CACNA2D3* involved in neurexin-mediated retrograde signaling (Tong *et al*. 2017) and may play an important role in pathways for learned vocalisation (Wada *et al*. 2004; Friedrich *et al*. 2019). Although the biological functions associated with these genes are broad, this result provides candidate genes for future studies investigating altered dispersion physiology and vocalisation.

The above discussed genes represent a short and analytically conservative list of genes putatively under selection across the starling’s global ranges. There are undoubtedly more sites of selection, either only identified in one of the outliers pairwise data sets (and so not categorised as parallel or divergent), in linkage with the identified SNPs (Brodie *et al*. 2016), or not sequenced at all using this reduced representation approach. However, these variants and genes serve as a shortlist of suitable targets for future gene expression analysis, or as the basis of further hypothesis on global avian selection pressures.

### 4.4 Historic sample sequencing

The success rate of historic sample sequencing was around 70%, with no correlation between sample properties and sequencing success. Patterns of density of variants across the genome identified using DArTseq appeared to follow similar patterns to the high-quality variant density data set provided by the whole genome comparison (Fig. 5), although historic samples did report a patchier variant distribution. However, prior simulations using data from historical samples reported that though historic samples contain significantly more missing data when compared to fresh tissue samples, the level of genotypic error had a minimal effect on population structure inference (Ewart *et al*. 2019).

In terms of sequence data processing, the BWA Aln-Stacks and BWA Mem-Stacks pipelines performed similarly, with bowtie-GATK performing comparatively much worse. This large difference may be due to the large amounts of missing data (Catchen *et al*. 2013), but are different from the relative performance previously reported for these two variant calling pipelines (Wright *et al*. 2019), suggesting that variant calling success is very data set dependent.

### 4.5 Future directions

Greater coverage of native range starling genetics is a vital future step for evolutionary genomic studies on this species. Improved native range genetic data will both help us better understand population structure and allelic shifts in the invasive ranges, and also shed light on native range dispersion dynamics. Further, the success of the museum sample sequencing demonstrated here gives hope that when largescale genomic studies are conducted in the native range, historical samples will be able to provide crucial background information regarding genetic diversity prior to the significant population declines.

Analyses of a greater number of historical samples will also aid in the categorisation of adaptive SNPs, as this will reduce possible effects of random sampling bias and capture more rare alleles. The sequencing failure rate of this study is comparable to another study using similarly aged museum skins (Ewart *et al*. 2019), suggesting that future projects seeking to use museum samples might expect similar failure rates (30-40%) and adjust their sampling design accordingly. Finally, similar analyses may be conducted between the historic UK and the well-studied contemporary North American starlings (as this population has a similar introduction time, range size, and environmental variation: Bodt *et al*. 2020; Hofmeister *et al*. 2021b), and then further extended upon through comparisons to other invasive populations in New Zealand, South Africa, and South America. Comparisons to parallel invasive populations will provide an invaluable opportunity to contrast concurrent species invasion and selection across multiple different continents, allowing for the discovery of broad evolutionary pattern in this invasive species.

## 5. Conclusions

Overall, this study demonstrates that the combination of native, invasive, and historical genetic data can lead to a more thorough understanding of global species shifts during the Anthropocene. We use genetic sequencing of museum specimens to identify putatively adaptive genetic changes through reduced representation sequencing and outlier SNP identification analysis. We have described evidence of parallel and divergent evolution in native and invasive starlings since the mid-19^th^ Century. Finally, we identify an apparent bias towards putatively adaptive SNPs on the Z chromosome, suggesting that the major sex chromosome may play an overly proportionate role in rapid evolution within this species.

## Acknowledgements

Thank you to Hein Van Grouw at Tring NHM for assistance with sourcing historical starling specimens. LAR was supported by a Scientia Fellowship from UNSW.

## Author Contributions

Project conception: KCS, LAR

Sample Collection: MB, ME, MCM Lab Work: KCS, JJA

Data Analysis: KCS Manuscript Writing: KCS

Manuscript Editing: All authors

## Data Accessibility

The data have been deposited with links to BioProject accession number XXXXX in the NCBI BioProject database (https://www.ncbi.nlm.nih.gov/bioproject/). Any scripts or metadata not covered by the above will be available on GitHub.

