## Supplementary Materials for "Using historical museum samples to examine divergent and parallel evolution in the invasive starling"

### Supplementary Material: Appendix 1

*Alternate variant calling pipelines*

In addition to the BWA *aln* pipeline, the BWA *mem* and GATK variant calling pipeline was run on the cleaned and processed raw data produced by *process_radtags*. BWA *mem* was run on default parameters, before being processed by STACKS *gstacks* and *populations*. For the GATK pipeline, BOWTIE2 was used for alignment (--phred33 --very-sensitive-local –I), SAMTOOLS to produce a sorted bam file. The PICARD v2.18.26 (under Java v8u121) *BuildBamIndex* function was used to index the reads. The GATK *HaplotypeCaller* function was used to call SNPs and assemble the haplotypes separately for eachs ample. The GATK functions *CombineGVCFs* and *GenotypeGVCFs* were used to combine each individual gvcf file into one vcf file for all individuals

For comparison to the primary variant data set, two filtering parameters were used. For BWA *mem*, no filtering parameters were used in *populations*, and a filtered file was produced using the exact same filtering parameters as for the primary variant calling pipeline pipeline (STACKS *populations*: -r 0.5 -p 2 --lnl_lim -15 --write_random_snp, and VCFTOOLS --max-missing 0.85 --maf 0.025 --minDP 5 --minGQ 15). For GATK, as the STACKS *population* function is not used, the filtering parameters could not be replicated exact, instead the VCFTOOLS parameters --max-missing 0.5 --maf 0.025 were used.

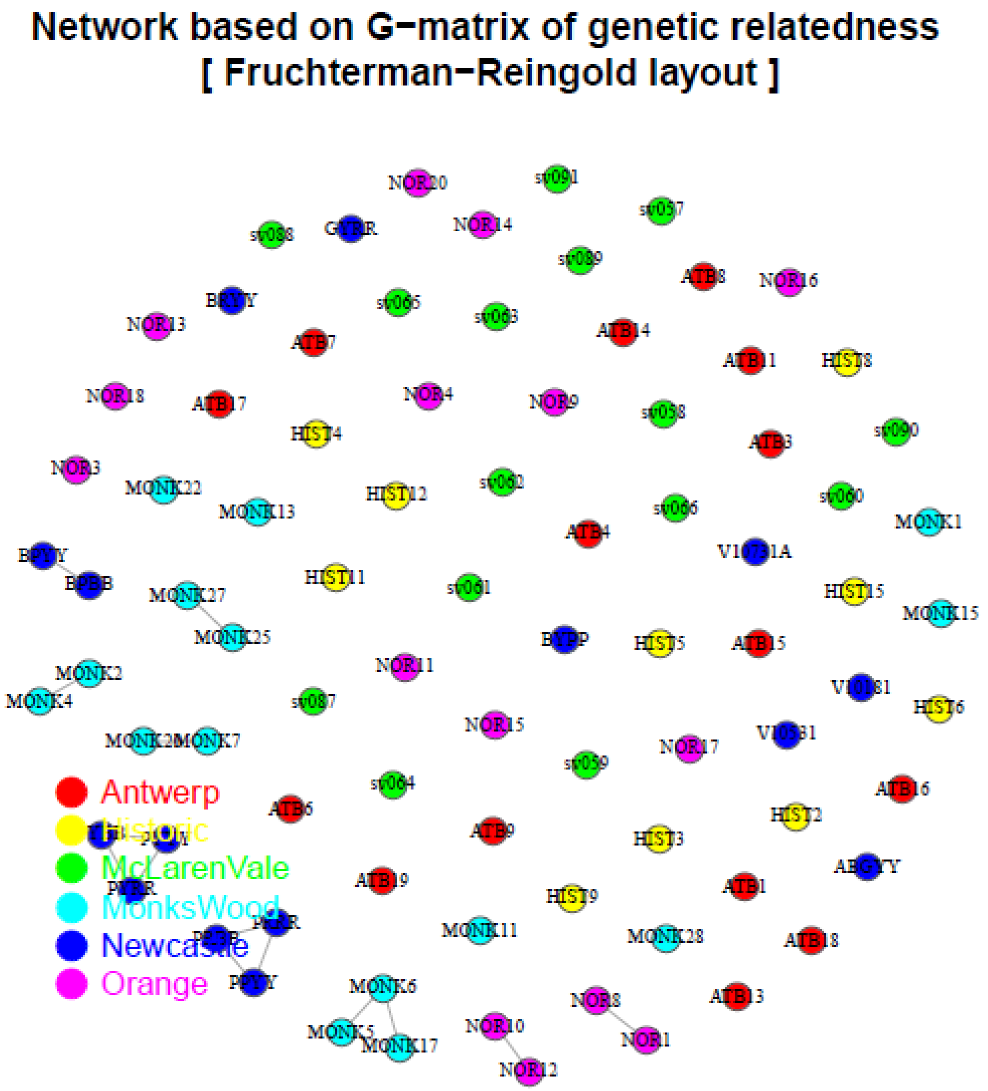

**Figure S1: Relatedness of *Sturnus vulgaris* contemporary and historical samples** using all SNP data, filtered, maf filter as well (hwe not excluded), with relatedness linkage displayed for individuals with a quantile of 0.004. Using the VCFTOOLS --relatedness2 flag reported these individuals above a relatedness threshold of >=0.2.

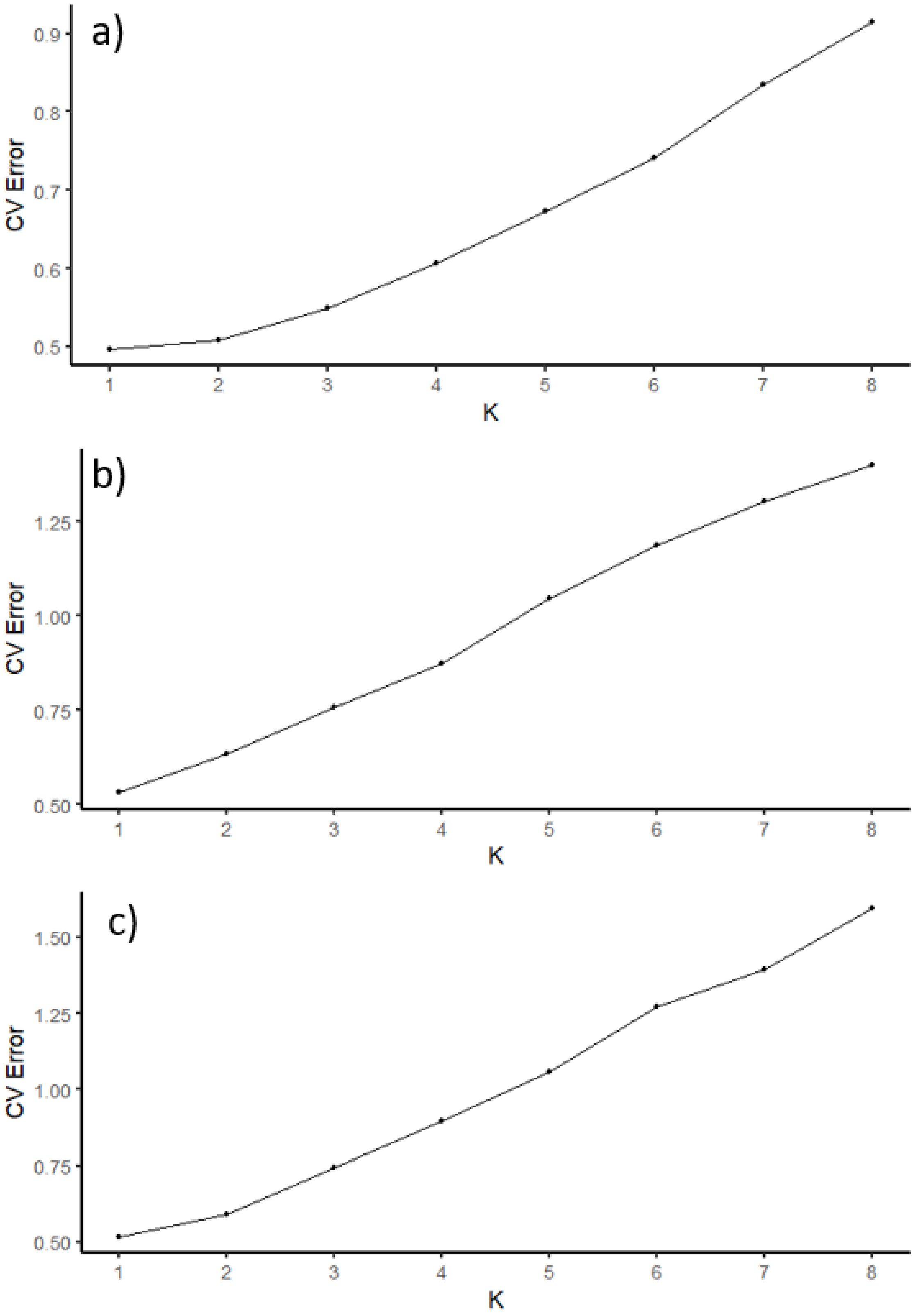

**Figure S2:** ***Sturnus vulgaris*** **admixture cross validation (CV) error profiles** for ADMIXTURE runs for a) all filtered data, b) UK + hist, and c) AU.

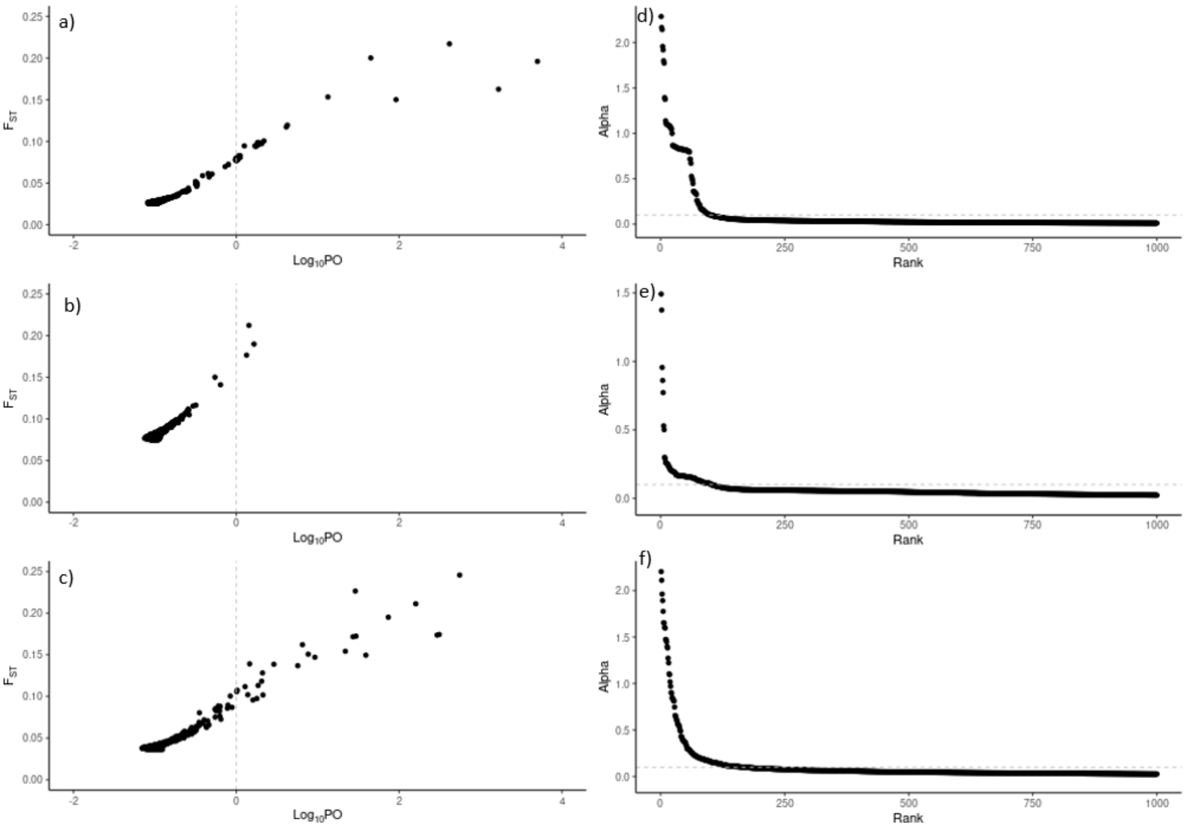

**Figure S3:** ***Sturnus vulgaris*** **outlier analysis BAYESCAN plots** of posterior odds log likelihood for pairwise data sets of a) UK-HS and c) AU-HS and e) UK-AU, and rank order plots for the top 1000 alpha values for b) UK-HS and d) AU-HS and f) UK-AU.

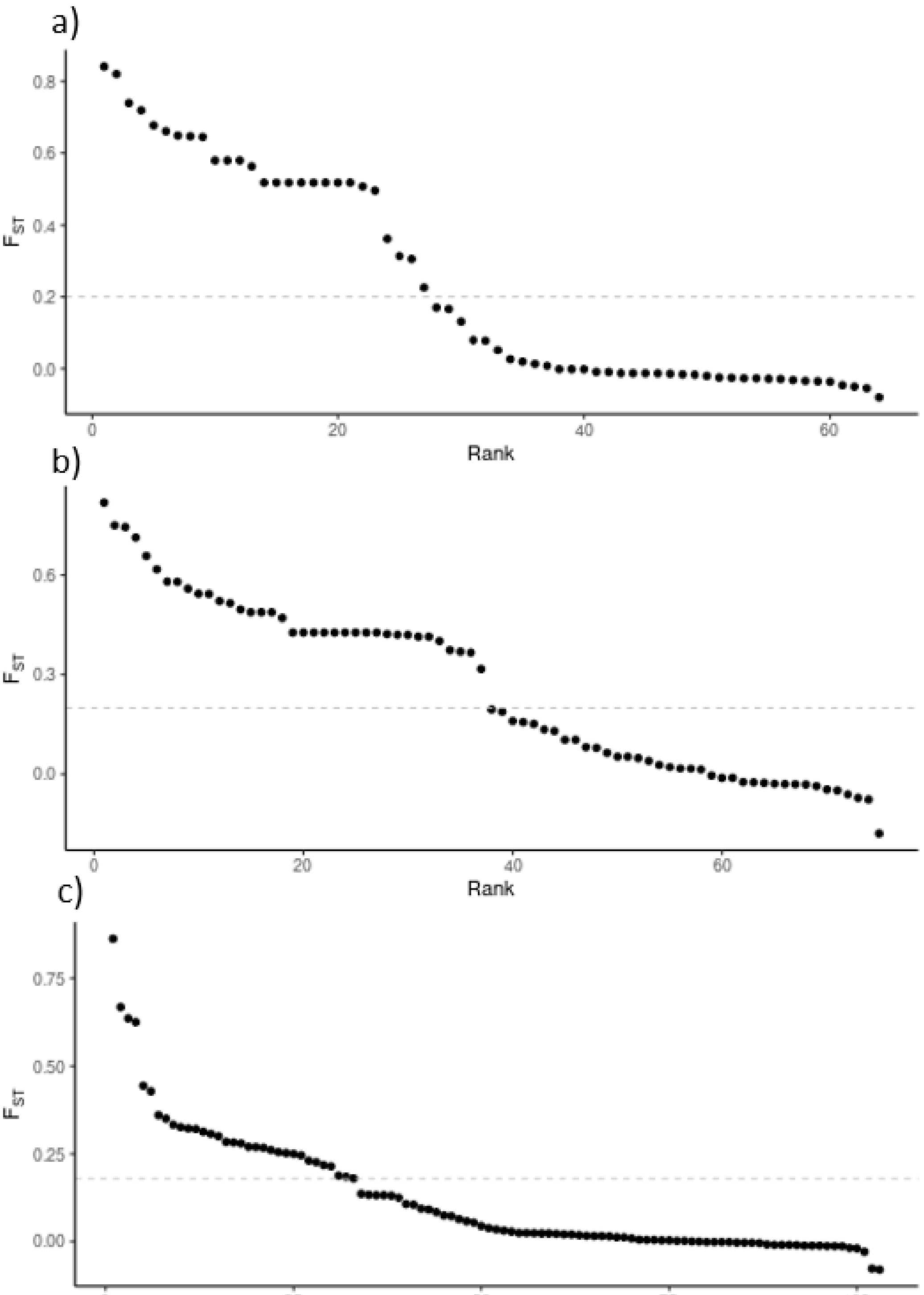

**Figure S4: *Sturnus vulgaris* outlier analysis SNP FST plots** for those flagged in the outlier sliding windows for pairwise data sets of a) UK-HS and b) AU-HS and c) UK-AU. SNPs left of vertical line were retained as outliers.

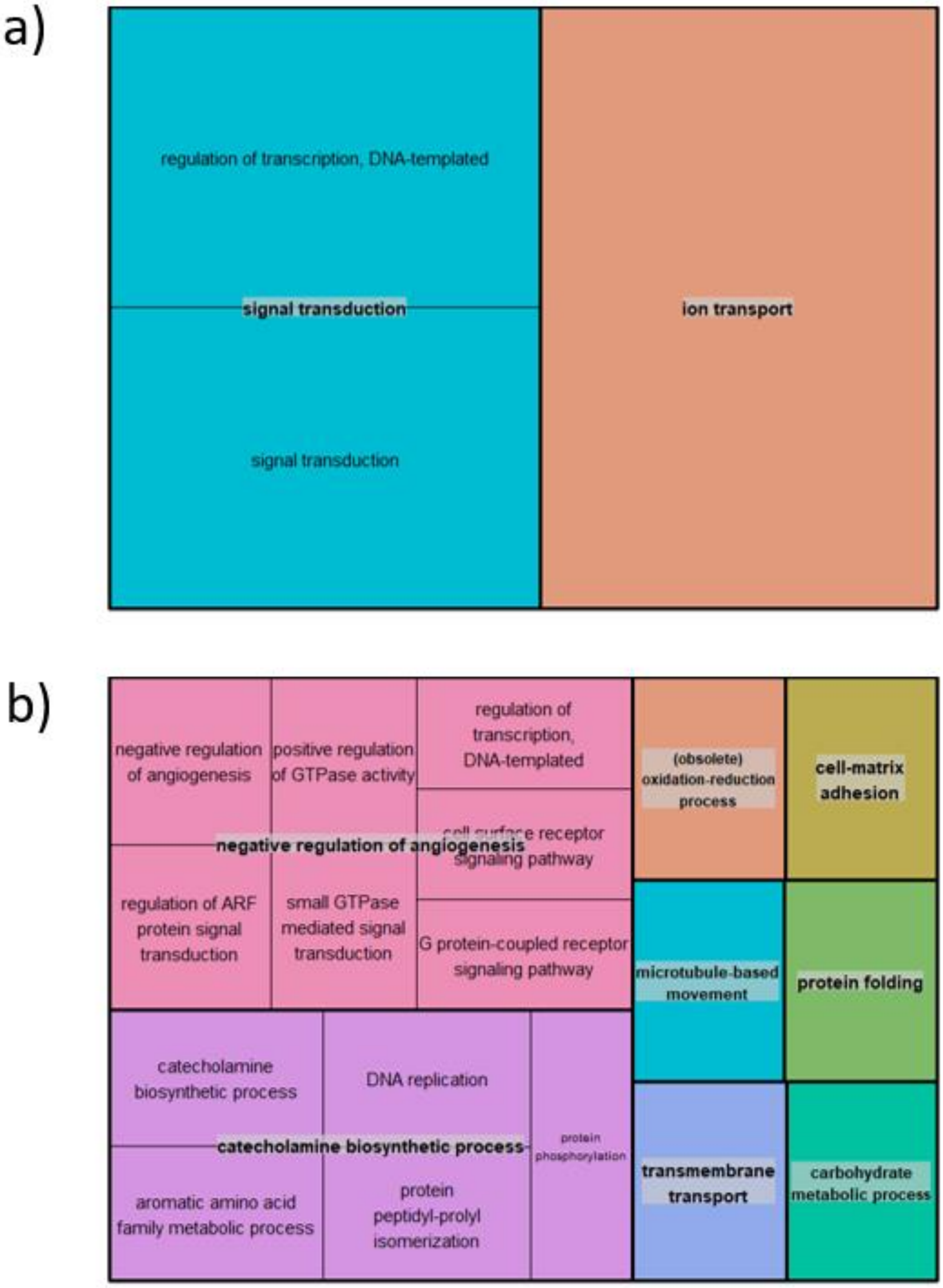

**Figure S5: REVIGO output of biological processes GO terms associated with putative genes under selection in *Sturnus vulgaris*** for SNPs reported in the a) divergent selection, and b) parallel selection data sets.

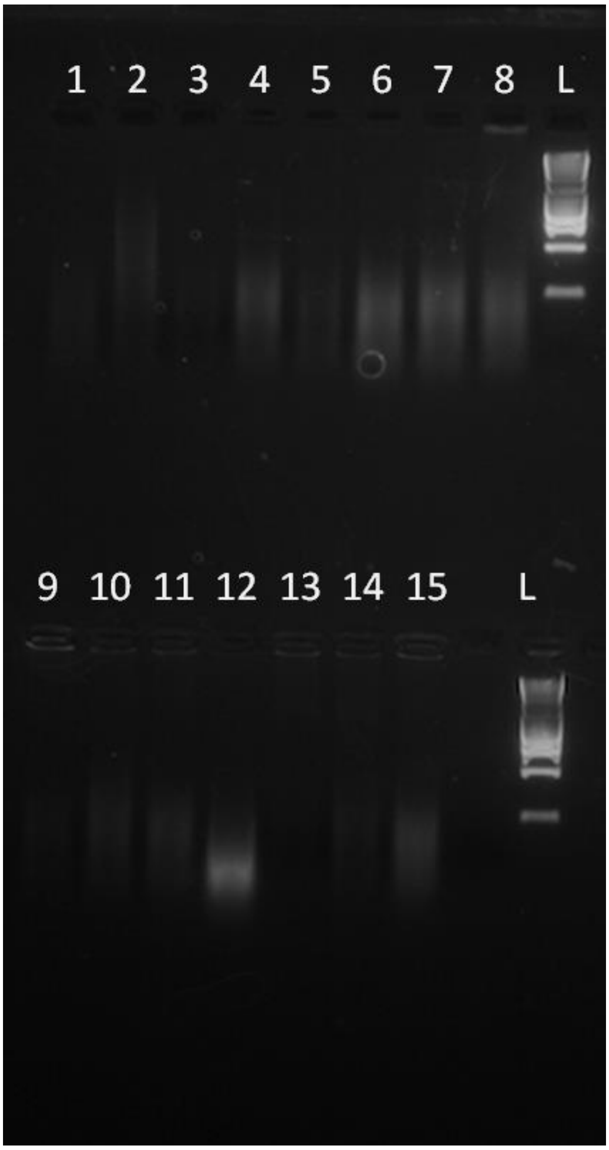

**Figure S6: Gel of historical *Sturnus vulgaris* samples**. 4ul of extracted DNA on 2% agrose gel. Numbers above wells denote sample ID corresponding to Table S1. L denotes HyperLadder1 (First band in HL I = 200bp, second band = 400 bp). Samples were diluted by approximately half from the concentrations reported in Table S1.

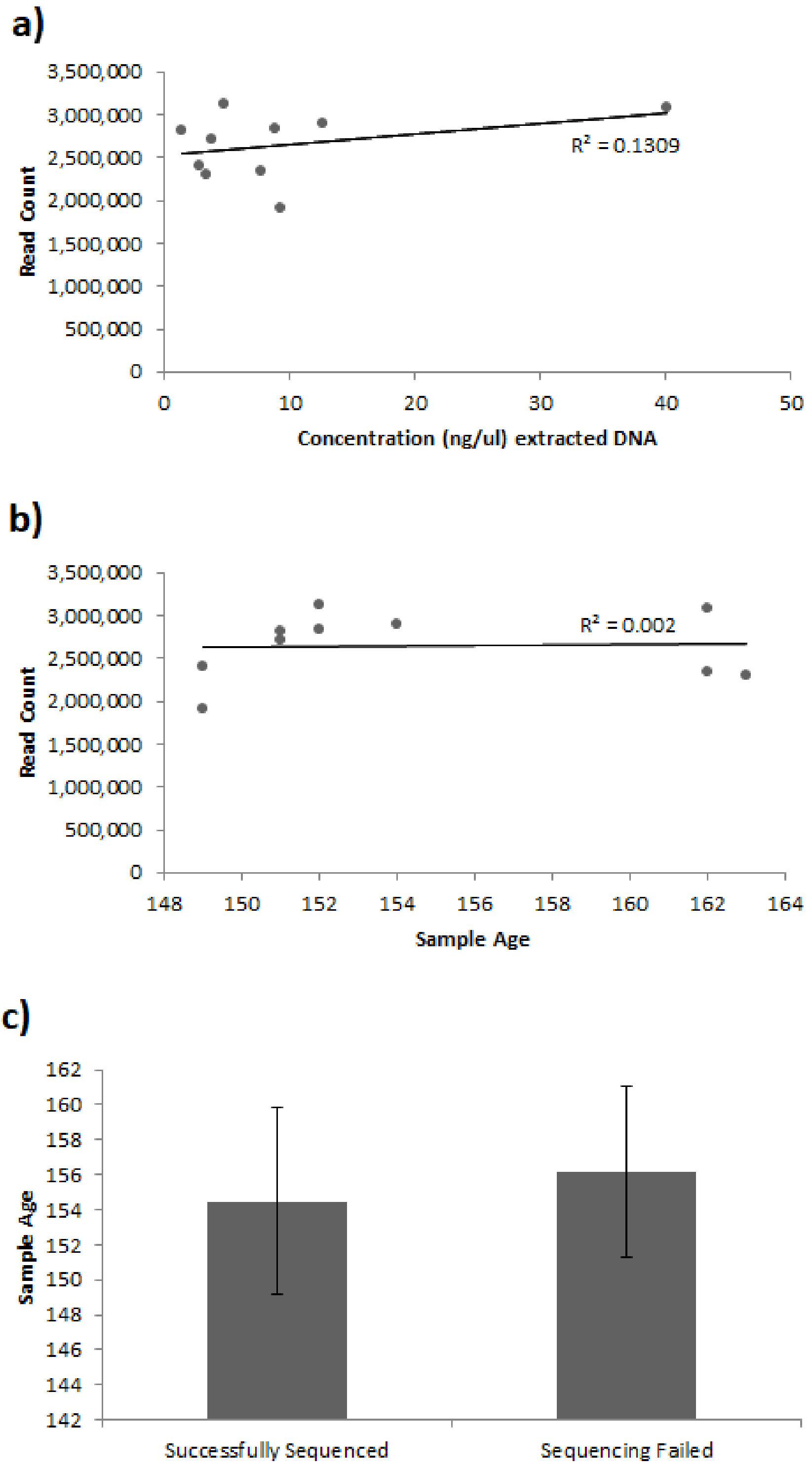

**Figure S7: Historical *Sturnus vulgaris* sample sequencing assessment** with panel a) depicting the relationship between raw read count and sample concentration, panel b) depicting the relationship between raw read count and sample age, and panel c) depicting the average sample age (+/- standard deviation) for the 10 successfully sequenced, and 5 failed historical samples.

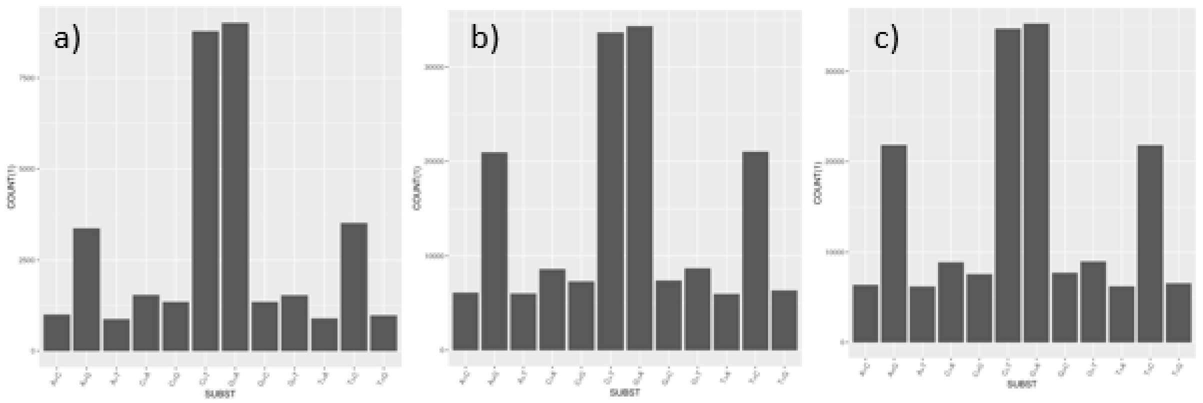

**Figure S8: *S vulgaris* SNP data per base substitution counts** in the unfiltered data set for a) historical, b) contemporary native range, and c) Australian *Sturnus vulgaris* samples.

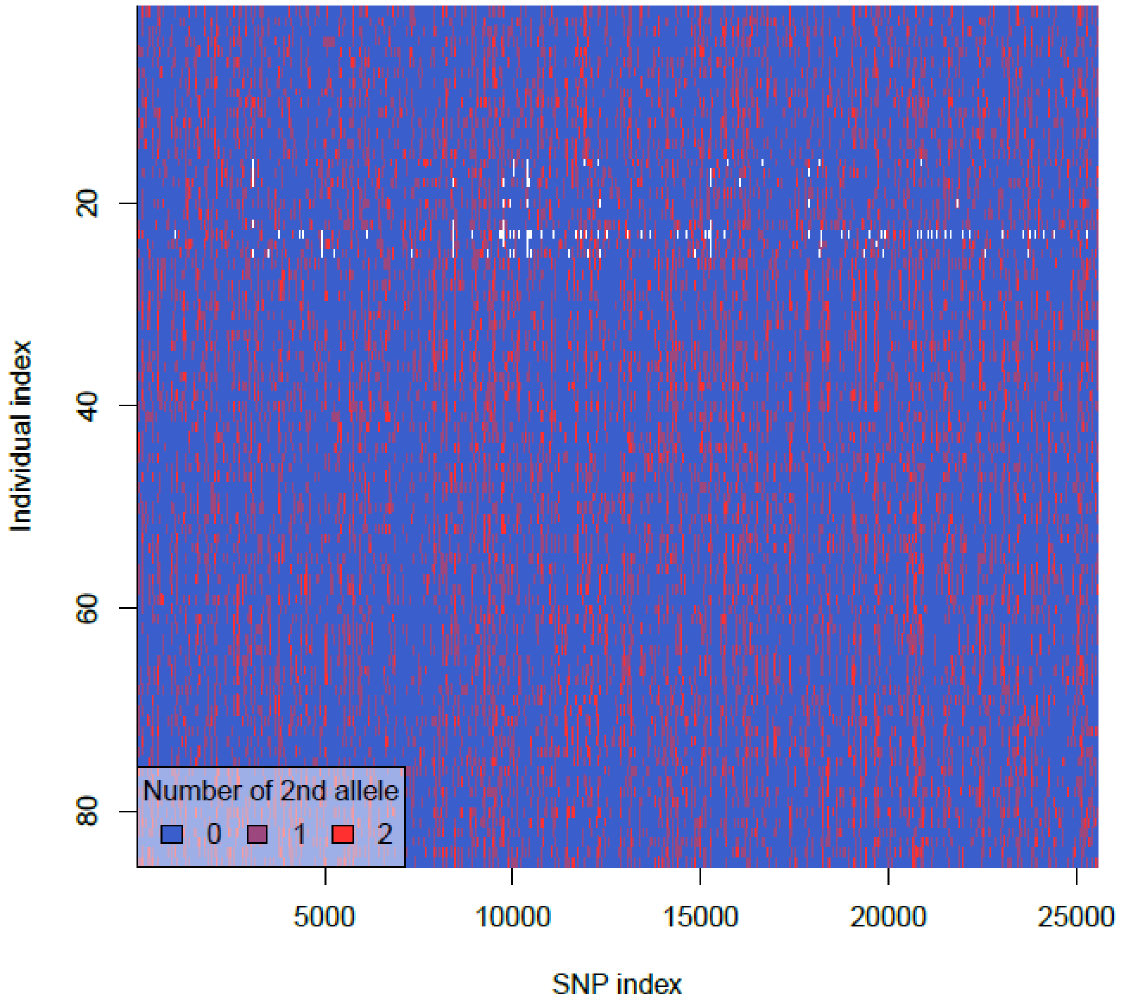

**Figure S9: Smear plot of *Sturnus vulgaris* reduced representation sequencing SNP data,** using the processed and filtered BWA-Aln variants.

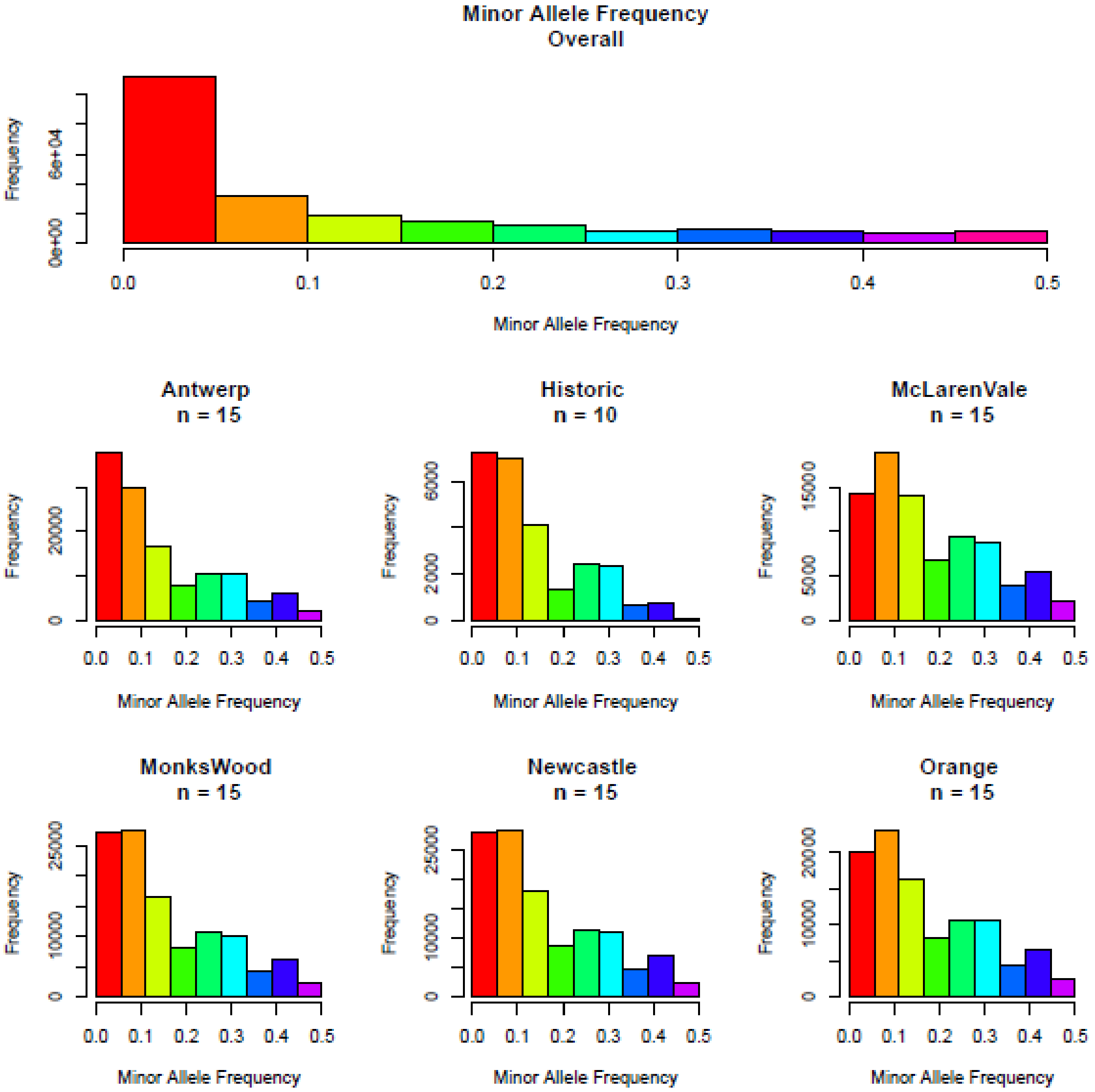

**Figure S10: *Sturnus vulgaris* reduced representation sequencing SNP data MAF profiles** for complete data set overall and per sample site.

**Table S1: *Sturnus vulgaris* historical native range museum samples (Tring NHM) metadata.**

| Sample ID | NHM ID | Date of Collection | Conc. (ng/ul) extracted DNA | Successfully sequenced | Reads Count |
|---|---|---|---|---|---|
| 1 | 87.7.1.2 | 01/1868 | 2.49 | NO | - |
| 2 | 2016.14.2 | 20/05/1869 | 3.82 | YES | 2,705,845 |
| 3 | 87.7.1.3 | 27/12/1869 | 1.40 | YES | 2,809,003 |
| 4 | 88.9.20.3 | 03/09/1868 | 4.78 | YES | 3,122,538 |
| 5 | 87.7.1.4 | 01/2/1871 | 2.86 | YES | 2,397,626 |
| 6 | 72.10.3.11 | 10/1871 | 9.34 | YES | 1,906,213 |
| 7 | 89.3.6.214 | 25/10/1866 | 10.18 | NO | - |
| 8 | 1889.3.6(215) | 17/03/1866 | 12.67 | YES | 2,902,553 |
| 9 | 79.4.5.879 | 05/1857 | 3.40 | YES | 2,295,671 |
| 10 | 97.11.10.729 | 22/06/1869 | 5.94 | NO | - |
| 11 | 81.5.1.3222 | 12/05/1858 | 7.81 | YES | 2,349,968 |
| 12 | 81.5.1.3227 | 05/06/1858 | 40.20 | YES | 3,076,038 |
| 13 | 81.5.1.3231 | 23/02/1857 | 1.72 | NO | - |
| 14 | 81.5.1.3232 | 31/1/1859 | 4.16 | NO | - |
| 15 | 81.5.3233 | 04/1868 | 8.85 | YES | 2,843,194 |

**Table S2: Mapping reads of historical and contemporary *Sturnus vulgaris*** using three different mapping pipelines; BWA-aln, BWA-mem, and Bowtie2-GATK. Mapped reads percentages were identified using the SAMTOOLS flagstat function. Loci and variant counts are as reported by either STACKS *populations* for the BWA mapping, or by SAMTOOLS for the GATK mapping. Individual missingness was calculated by using the VCFTOOLS --missing-indv flag.

| | | **Raw Data** | **Unfiltered data set** | | **Filtered data set** | |
|---|---|---|---|---|---|---|
| | | **Mapped reads %** | **No. of loci/variant sites** | **Average Ind Data Missing (%)** | **No. of variant sites** | **Average Ind Data Missing (%)** |
| **BWAAln-Stacks** | **Contemporary & Historical** | 57.63 | 411,298/239,538 | 35.85 (historic: 83.28) | 13,722 | 10.08 (historic: 80.43) |
| | **Historical** | 8.53 | 250,463/30,662 | 35.07 | 2,744 | 5.00 |
| **BWAMem-stacks** | **Contemporary & Historical** | 85.36 | 412,079/243,589 | 34.76 (historic: 81.57) | 14,929 | 9.61 (historic: 76.63) |
| | **Historical** | 53.95 | 263,686/34,832 | 33.05 | 4,331 | 4.13 |
| **Bowtie2-GATK** | **Contemporary & Historical** | 89.17 | 5879/5015 | 60.25 (historic: 74.58) | 715 | 28.61 (51.03) |
| | **Historical** | 65.24 | 1235/791 | 57.51 | 500 | 25.08 |

**Table S3:** ***Sturnus vulgaris*** **SNPs discovered by each outlier identification method.** Diagonal is unique to method. Overlap indicates a SNPs presence in present in both approaches. Total number of SNPs present across all 3 identification methods are listed in each of the pairwise population subheadings.

| **UK-HIST (4 SNPs present across all)** | | | |
|---|---|---|---|
| | **Bayescan FDR (8)** | **Bayescan alpha 005 + LD (62)** | **Windows 0.99 (27)** |
| **Bayescan FDR (8)** | 2 | - | - |
| **Bayescan alpha 01 + LD (62)** | 6 | 40 | - |
| **Windows 0.99 (27)** | 4 | 20 | 7 |
| **AU-HIST (1 SNPs present across all)** | | | |
| | **Bayescan FDR (1)** | **Bayescan alpha 005 + LD (69)** | **Windows 0.99 (37)** |
| **Bayescan FDR (1)** | 0 | - | - |
| **Bayescan alpha 01 + LD (69)** | 1 | 48 | - |
| **Windows 0.99 (37)** | 1 | 21 | 16 |
| **UK-AU (3 SNPs present across all)** | | | |
| | **Bayescan FDR (15)** | **Bayescan alpha 005 + LD (34)** | **Windows 0.99 (33)** |
| **Bayescan FDR (15)** | 10 | - | - |
| **Bayescan alpha 01 + LD (34)** | 4 | 27 | - |
| **Windows 0.99 (33)** | 4 | 6 | 26 |

**Table S4: Summary of allelic frequency for outlier SNPs in *Sturnus vulgaris*** under divergent and parallel selection as calculated by STACKS.

| CLASSIFY | CHROM | POS | Mapped Gene/Genes | Major Allele | UK Major freq | AU Major freq | HIST Major freq |
|---|---|---|---|---|---|---|---|
| Divergent | starling4 | 1.26E+08 | | A | 0.965517 | 0.625 | 1 |
| Divergent | starling5 | 23635575 | | G | 0.571429 | 0.166667 | 0.875 |
| Divergent | starling5 | 24369993 | GRIK2 | G | 0.75 | 0 | 0.75 |
| Divergent | starling5 | 67026996 | Esrrg | C | 0.361111 | 0.839286 | 0.25 |
| Divergent | starling6 | 57107109 | ARHGAP10 | C | 0.68 | 1 | 0.3 |
| Divergent | starling15 | 4462195 | Cacna2d3 | C | 1 | 0.461538 | 1 |
| Divergent | starling16 | 18573988 | ANKHD1 | G | 0.955556 | 0.666667 | 0.75 |
| Divergent | starling26 | 5999859 | | A | 1 | 0.6 | 0.6 |
| Divergent | starling30 | 4287460 | Unknown protein | C | 0.142857 | 1 | 1 |
| Parallel | starling2 | 3428749 | | T | 1 | 1 | 0.666667 |
| Parallel | starling2 | 35585267 | GRM5 | C | 0.944444 | 1 | 0.5 |
| Parallel | starling2 | 42488998 | | A | 0.96 | 1 | 0.611111 |
| Parallel | starling2 | 71327811 | | C | 1 | 1 | 0.666667 |
| Parallel | starling2 | 72180065 | C2CD2 | A | 1 | 1 | 0.666667 |
| Parallel | starling2 | 83513325 | | G | 0.90625 | 0.928571 | 0.357143 |
| Parallel | starling3 | 12791521 | Orc5 | G | 1 | 1 | 0.7 |
| Parallel | starling3 | 14914566 | SI | G | 1 | 1 | 0.7 |
| Parallel | starling3 | 29905895 | | G | 1 | 1 | 0.7 |
| Parallel | starling3 | 51874566 | Unknown protein | G | 1 | 1 | 0.7 |
| Parallel | starling4 | 7802066 | | G | 1 | 1 | 0.714286 |
| Parallel | starling4 | 27963762 | | G | 1 | 1 | 0.7 |
| Parallel | starling4 | 42474388 | | G | 1 | 1 | 0.7 |
| Parallel | starling4 | 46658977 | | C | 1 | 1 | 0.7 |
| Parallel | starling4 | 58362459 | | T | 1 | 1 | 0.666667 |
| Parallel | starling4 | 64264491 | PIEZO2 | A | 1 | 1 | 0.666667 |
| Parallel | starling4 | 70917693 | DNAH5 | C | 1 | 1 | 0.6 |
| Parallel | starling4 | 71666440 | GABBR2 | C | 1 | 1 | 0.7 |
| Parallel | starling4 | 99568041 | Unknown protein | G | 0.97561 | 0.948276 | 0.35 |
| Parallel | starling4 | 1.24E+08 | | C | 1 | 1 | 0.7 |
| Parallel | starling5 | 6572134 | | A | 1 | 1 | 0.666667 |
| Parallel | starling5 | 37412510 | ADGRB3 | G | 1 | 1 | 0.666667 |
| Parallel | starling5 | 41375430 | SOS1 | C | 1 | 1 | 0.7 |

| | | | | | | | |
|---|---|---|---|---|---|---|---|
| Parallel | starling5 | 43588236 | | G | 1 | 1 | 0.7 |
| Parallel | starling5 | 51303448 | | G | 1 | 1 | 0.7 |
| Parallel | starling5 | 56953867 | SAMD3 | G | 1 | 1 | 0.75 |
| Parallel | starling5 | 71978252 | | G | 0.5 | 0.533333 | 1 |
| Parallel | starling5 | 1.03E+08 | INS | C | 1 | 1 | 0.7 |
| Parallel | starling7 | 20803249 | | C | 1 | 1 | 0.7 |
| Parallel | starling7 | 21203978 | TRIP4 | C | 1 | 1 | 0.666667 |
| Parallel | starling8 | 10152279 | SBF2 | C | 1 | 1 | 0.7 |
| Parallel | starling8 | 39002654 | | C | 1 | 1 | 0.7 |
| Parallel | starling8 | 52016888 | SNX30/SLC46A2 | G | 1 | 1 | 0.666667 |
| Parallel | starling8 | 56030454 | NID2 | T | 1 | 1 | 0.7 |
| Parallel | starling10 | 28352666 | | C | 0.977778 | 1 | 0.625 |
| Parallel | starling13 | 14681104 | | C | 1 | 1 | 0.7 |
| Parallel | starling14 | 18202987 | | C | 1 | 1 | 0.666667 |
| Parallel | starling15 | 9046251 | PXK | G | 0.2 | 0.454545 | 1 |
| Parallel | starling15 | 19435080 | IQSEC1 | G | 1 | 1 | 0.7 |
| Parallel | starling16 | 3396982 | Trim25 | G | 1 | 1 | 0.666667 |
| Parallel | starling19 | 2497428 | Unknown protein | G | 1 | 1 | 0.7 |
| Parallel | starling20 | 12083219 | ACSF2/Chad | G | 1 | 1 | 0.7 |
| Parallel | starling21 | 8454504 | mybbp1a | C | 0.939394 | 0.913793 | 0.25 |
| Parallel | starling23 | 8054190 | DDX19A | C | 1 | 1 | 0.7 |
| Parallel | starling31 | 7189771 | UBAP2 | C | 1 | 1 | 0.7 |
| Parallel | starling31 | 10663913 | IL7R | A | 1 | 1 | 0.7 |
| Parallel | starling31 | 18108153 | | T | 1 | 1 | 0.7 |
| Parallel | starling31 | 19762165 | CWC27 | C | 1 | 1 | 0.666667 |
| Parallel | starling31 | 21621921 | TENT2 | G | 0.947368 | 0.96 | 0.285714 |
| Parallel | starling31 | 24764608 | | A | 0.966667 | 0.95 | 0.142857 |
| Parallel | starling31 | 33855811 | | C | 1 | 1 | 0.625 |
| Parallel | starling31 | 52606417 | Pak3 | G | 1 | 1 | 0.25 |

**Table S5: Summary of putative genes under spatial and temporal selection in *Sturnus vulgaris*** that were mapped to by the outlier SNPs reported at putatively divergent or parallel. Definitions pulled from GeneCards (Stelzer *et al.* 2016).

| Gene | Name | Biological Function |
|---|---|---|
| **Divergent Genes** | | |
| *GRIK2* | Glutamate Ionotropic Receptor Kainate Type Subunit 2 | Glutamate receptors are the predominant excitatory neurotransmitter receptors in the mammalian brain and are activated in a variety of normal neurophysiologic processes. Mutations in this gene have been associated with autosomal recessive cognitive disability. |
| Esrrg | estrogen related receptor gamma | This gene encodes a member of the estrogen receptor-related receptor (ESRR) family, which belongs to the nuclear hormone receptor superfamily. It has been reported that the family member encoded by this gene functions as a transcriptional activator of DNA cytosine-5-methyltransferases 1 (Dnmt1) expression by direct binding to its response elements in the DNMT1 promoters, modulates cell proliferation and estrogen signaling in breast cancer, and negatively regulates bone morphogenetic protein 2-induced osteoblast differentiation and bone formation. |
| ARHGAP10 | Rho GTPase-activating protein 10 | GTPase activator for the small GTPases RhoA and Cdc42 by converting them to an inactive GDP-bound state. Essential for PTKB2 regulation of cytoskeletal organization via Rho family GTPases. |
| Cacna2d3 | Voltage-dependent calcium channel subunit alpha-2/delta-3 | The alpha-2/delta subunit of voltage-dependent calcium channels regulates calcium current density and activation/inactivation kinetics of the calcium channel. |
| ANKHD1 | Ankyrin repeat and KH domain-containing protein 1 | May play a role as a scaffolding protein that may be associated with the abnormal phenotype of leukemia cells. Isoform 2 may possess an antiapoptotic effect and protect cells during normal cell survival through its regulation of caspases. |
| **Parallel Genes** | | |
| GRM5 | Metabotropic glutamate receptor 5 | Is a metabatropic glutamate receptor, whose signaling activates a phosphatidylinositol-calcium second messenger system. This protein may be involved in the regulation of neural network activity and synaptic plasticity. Glutamatergic neurotransmission is involved in most aspects of normal brain function and can be perturbed in many neuropathologic conditions. |
| C2CD2 | C2 domain-containing protein 2 | Lipid-binding protein that transports phosphatidylinositol, the precursor of phosphatidylinositol 4,5-bisphosphate (PI(4,5)P2), from its site of synthesis in the endoplasmic reticulum to the cell membrane. |

| | | |
|---|---|---|
| Orc5 | Origin recognition complex subunit 5 | The origin recognition complex (ORC) is a highly conserved six subunit protein complex essential for the initiation of the DNA replication in eukaryotic cells. |
| SI | Sucrase-Isomaltase, Intestinal | This gene encodes a sucrase-isomaltase enzyme that is expressed in the intestinal brush border. The encoded protein is synthesized as a precursor protein that is cleaved by pancreatic proteases into two enzymatic subunits sucrase and isomaltase. These two subunits heterodimerize to form the sucrose-isomaltase complex. This complex is essential for the digestion of dietary carbohydrates including starch, sucrose and isomaltose. |
| PIEZO2 | Piezo-type mechanosensitive ion channel component 2 | The protein encoded by this gene contains more than thirty transmembrane domains and likely functions as part of mechanically-activated (MA) cation channels. These channels serve to connect mechanical forces to biological signals. The encoded protein quickly adapts MA currents in somatosensory neurons. |
| DNAH5 | Dynein Axonemal Heavy Chain 5 | This gene encodes a dynein protein, which is part of a microtubule-associated motor protein complex consisting of heavy, light, and intermediate chains. This protein is an axonemal heavy chain dynein. It functions as a force-generating protein with ATPase activity, whereby the release of ADP is thought to produce the force-producing power stroke. |
| GABBR2 | Gamma-aminobutyric acid type B receptor subunit 2 | The multi-pass membrane protein encoded by this gene belongs to the G-protein coupled receptor 3 family and GABA-B receptor subfamily. The GABA-B receptors inhibit neuronal activity through G protein-coupled second-messenger systems, which regulate the release of neurotransmitters, and the activity of ion channels and adenylyl cyclase. Allelic variants of this gene have been associated with nicotine dependence. |
| ADGRB3 | Adhesion G protein-coupled receptor B3 | This p53-target gene encodes a brain-specific angiogenesis inhibitor, a seven-span transmembrane protein, and is thought to be a member of the secretin receptor family, and may also play a role in angiogenesis. |
| SOS1 | Son of sevenless homolog 1 | This gene encodes a protein that is a guanine nucleotide exchange factor for RAS proteins, membrane proteins that bind guanine nucleotides and participate in signal transduction pathways. GTP binding activates and GTP hydrolysis inactivates RAS proteins. The product of this gene may regulate RAS proteins by facilitating the exchange of GTP for GDP. Mutations in this gene are associated with gingival fibromatosis 1 and Noonan syndrome type 4. |
| SAMD3 | Sterile alpha motif domain-containing protein 3 | This gene encodes a multidomain protein that functions as a scaffold protein to mediate the mitogen-activated protein kinase pathways downstream from Ras. This gene product is induced by vitamin D and inhibits apoptosis in certain cancer cells. It may also play a role in ternary complex assembly of synaptic proteins at the postsynaptic membrane and coupling of signal transduction to membrane/cytoskeletal remodeling. |
| INS | Insulin | This gene encodes insulin, a peptide hormone that plays a vital role in the regulation of carbohydrate and lipid metabolism. Binding of insulin to the insulin receptor (INSR) stimulates glucose uptake. |
| TH | Tyrosine 3-monooxygenase | The protein encoded by this gene is involved in the conversion of tyrosine to dopamine. It is the rate-limiting enzyme in the synthesis of catecholamines, hence plays a key role in the physiology of adrenergic neurons. |

| | | |
|---|---|---|
| TRIP4 | Activating signal cointegrator 1 | This gene encodes a subunit of the tetrameric nuclear activating signal cointegrator 1 (ASC-1) complex, which associates with transcriptional coactivators, nuclear receptors and basal transcription factors to facilitate nuclear receptors-mediated transcription. This protein is localized in the nucleus and contains an E1A-type zinc finger domain, which mediates interaction with transcriptional coactivators and ligand-bound nuclear receptors, such as thyroid hormone receptor and retinoid X receptor alpha, but not glucocorticoid receptor. |
| SBF2 | Myotubularin-related protein 13 | Promotes the exchange of GDP to GTP, converting inactive GDP-bound Rab proteins into their active GTP-bound form. |
| SNX30 | Sorting nexin-30 | This gene encodes a member of the sorting nexin family. Members of this family contain a phox (PX) domain, which is a phosphoinositide binding domain, and are involved in intracellular trafficking. |
| SLC46A2 | Thymic stromal cotransporter homolog | SLC46A2 (Solute Carrier Family 46 Member 2) is a Protein Coding gene. Gene Ontology (GO) annotations related to this gene include symporter activity. |
| NID2 | Nidogen-2 | This gene encodes a member of the nidogen family of basement membrane proteins. This protein is a cell-adhesion protein that binds collagens I and IV and laminin and may be involved in maintaining the structure of the basement membrane. |
| PXK | PX domain-containing protein kinase-like protein | This gene encodes a phox (PX) domain-containing protein which may be involved in synaptic transmission and the ligand-induced internalization and degradation of epidermal growth factors. |
| IQSEC1 | IQ motif and SEC7 domain-containing protein 1 | Gene Ontology (GO) annotations related to this gene include lipid binding and ARF guanyl-nucleotide exchange factor activity. |
| Trim25 | E3 ubiquitin/ISG15 ligase TRIM25 | The protein encoded by this gene is a member of the tripartite motif (TRIM) family. The TRIM motif includes three zinc-binding domains, a RING, a B-box type 1 and a B-box type 2, and a coiled-coil region. The protein localizes to the cytoplasm. The presence of potential DNA-binding and dimerization-transactivation domains suggests that this protein may act as a transcription factor, similar to several other members of the TRIM family. Expression of the gene is upregulated in response to estrogen. |
| Gpsm1 | G-protein-signaling modulator 1 | G-protein signaling modulators (GPSMs) play diverse functional roles through their interaction with G-protein subunits. This gene encodes a receptor-independent activator of G protein signaling, which is one of several factors that influence the basal activity of G-protein signaling systems. The protein contains seven tetratricopeptide repeats in its N-terminal half and four G-protein regulatory (GPR) motifs in its C-terminal half. |
| ACSF2 | Acyl-CoA Synthetase Family Member 2 | ACSF2 (Acyl-CoA Synthetase Family Member 2) is a Protein Coding gene. Among its related pathways are Metabolism and Fatty Acyl-CoA Biosynthesis. Gene Ontology (GO) annotations related to this gene include ligase activity. |
| Chad | Chondroadherin | Chondroadherin is a cartilage matrix protein thought to mediate adhesion of isolated chondrocytes. |
| mybbp1a | Myb-binding protein 1A-like protein | This gene encodes a nucleolar transcriptional regulator that was first identified by its ability to bind specifically to the Myb proto-oncogene protein. The encoded protein is thought to play a role in many cellular processes including response to nucleolar stress, tumor suppression and synthesis of ribosomal DNA. |

| | | |
|---|---|---|
| DDX19A | ATP-dependent RNA helicase DDX19A | DDX19A (DEAD-Box Helicase 19A) is a Protein Coding gene. Diseases associated with DDX19A include Heavy Chain Disease. Among its related pathways are mRNA surveillance pathway and RNA transport. Gene Ontology (GO) annotations related to this gene include nucleic acid binding and helicase activity. |
| UBAP2 | Ubiquitin-associated protein 2 | The protein encoded by this gene contains a UBA (ubiquitin associated) domain, which is characteristic of proteins that function in the ubiquitination pathway. |
| IL7R | Interleukin-7 receptor subunit alpha | This protein has been shown to play a critical role in V(D)J recombination during lymphocyte development. |
| CWC27 | CWC27 spliceosome associated cyclophilin | Gene Ontology (GO) annotations related to this gene include peptidyl-prolyl cis-trans isomerase activity. |
| TENT2 | Poly(A) RNA polymerase GLD2 | Cytoplasmic poly(A) RNA polymerase that adds successive AMP monomers to the 3'-end of specific RNAs, forming a poly(A) tail. |
| Pak3 | Serine/threonine-protein kinase PAK 3 | The protein encoded by this gene is a serine-threonine kinase and forms an activated complex with GTP-bound RAS-like (P21), CDC2 and RAC1. This protein may be necessary for dendritic development and for the rapid cytoskeletal reorganization in dendritic spines associated with synaptic plasticity. |
